# On the prediction of DNA-binding preferences of C2H2-ZF domains using structural models: application on human CTCF

**DOI:** 10.1101/859934

**Authors:** Alberto Meseguer, Filip Årman, Oriol Fornes, Ruben Molina, Jaume Bonet, Narcis Fernandez-Fuentes, Baldo Oliva

## Abstract

Cis2-His2 zinc finger (C2H2-ZF) proteins are the largest family of transcription factors in human and higher metazoans. However, the DNA-binding preferences of many members of this family remain unknown. We have developed a computational method to predict these DNA-binding preferences. We combine information from crystal structures composed by C2H2-ZF domains and from bacterial one-hybrid experiments to compute scores for protein-DNA binding based on statistical potentials. We apply the scores to compute theoretical position weight matrices (PWMs) of proteins with a DNA-binding domain composed by C2H2-ZF domains, with the only requirement of an input structure (experimentally determined or modelled). We have tested the capacity to predict PWMs of zinc finger domains, successfully predicting 3-2 nucleotides of a trinucleotide binding site for about 70% variants of single zinc-finger domains of Zif268. We have also tested the capacity to predict the PWMs of proteins composed by three C2H2-ZF domains, successfully matching between 60% and 90% of the binding-site motif according to the JASPAR database. The tests are used as a proof of the capacity to scan a DNA fragment and find the potential binding sites of transcription-factors formed by C2H2-ZF domains. As an example, we have tested the approach to predict the DNA-binding preferences of the human chromatin binding factor CTCF.

## INTRODUCTION

The identification of transcription factors (TFs) and the characterization of their binding sites is key to understand how gene expression is regulated. Experimental techniques such as ChIP^1^, PBM^2^, HT-SELEX^3^, MPRA^4^ or bacterial and yeast one-hybrid^5,6^ have allowed the characterization of TF-binding sites at large-scale. However, experimental techniques are expensive and time consuming, and yet the binding preferences of many TFs remain unknown^7,8^. Given the current limitations, the usage of computational tools to complement experimental techniques is necessary.

Cis2-His2 zinc finger (C2H2-ZF) proteins are the largest family of TFs in higher metazoans^9^. They represent around the 45% of all known human TFs, being the largest TF family in human TFs^8^. C2H2-ZF proteins are involved in a wide range of biological processes such as development^10^ or chromatin compartmentalization^11^. C2H2-ZF proteins have been related to many diseases^12,13^ and can be used as tools for precise gene editing^14,15^. At this point, knowing the binding preferences of C2H2-ZF proteins becomes crucial, despite for many are yet unknown^8^. Besides, many members of the C2H2-ZF do not have close homologs across metazoans and thus, sequence homology cannot be used to infer their binding preferences^16^. Still, all of them have the same structure in the DNA binding domain. DNA binding domains (DBD) of C2H2-ZF proteins are composed by small domains called zinc fingers arranged in tandem^17^. Each zinc finger is able to recognize DNA sequences of 3 nucleotides^18^ and, by combining adjacent zinc fingers, C2H2-ZF proteins are able to recognize long and complex DNA patterns^19^. Human C2H2-ZF proteins contain an average of around 10 domains, leading to binding sites of about 30 bases^20^.

Several computational tools have been developed to predict the binding preferences of transcription factors and in particular C2H2-ZF proteins. Some tools are based on combining experimental data with the structure of the interaction between proteins and DNA. Among them, some approaches use different machine learning algorithms: random forest regression^16^, support vector machines^21,22^, single layer perceptrons^23^, hidden Markov models^24^ and other statistical models^25,26^; using residue-residue contacts^16,21,22,26^ or context dependencies and sequence similarities^23,24,27,28^. Other tools are based on the analysis of the structural patterns extracted from protein-DNA interactions^25,29-32^, or from the flexibility on the DNA chain, of which recent studies show the relevance of the DNA shape to consider the DNA-binding of a protein^33^. Some of these structure-based tools use statistical potentials^31,32^. Statistical potentials (also known as knowledge-based potentials) are scoring functions derived from the analysis of contacts in a set of structures. Statistical potentials have been widely used to evaluate the quality and the stability of protein folds, protein-protein interactions and protein-DNA interactions^34^.

Here we present a computational tool to predict the binding preferences of C2H2-ZF proteins. We combine experimental bacterial one-hybrid (B1H) data with structural, three-dimensional, information of TF-DNA complexes. We use B1H data^19^ to model TF-DNA interactions and increase the statistic power of the potentials by using computationally derived structural models for TF-DNA complexes with unknown structure. We use the potentials to predict the Position-Weight-Matrix (PWM) of the binding-site of C2H2-ZF domains^35,36^. We also predict PWMs for transcription factor proteins with three C2H2-ZF domains and compare them with their motifs in the JASPAR database ^37^. Finally, we apply statistical potentials to predict the binding preferences of human CTCF, a transcriptional repressor with a key role in genome compartmentalization^11^.

## METHODS

### 1. Software

The following software was used in this study: DSSP (version CMBI 2006) ^38^ to obtain protein structural features; X3DNA (version 2.0) ^39^ to analyze and generate DNA structures; *matcher* and *needle*, from the EMBOSS package (version 6.5.0) ^40^, to obtain local and global alignments, respectively; BLAST (version 2.2.22) ^41^ to search homologs of a given query (target) protein sequence; MODELLER (version 9.9) ^42^ to construct structural models; and the programs FIMO and TOMTOM from the MEME suite^43^ to scan a DNA sequence with a Position-Weight Matrix and to compare two PWMs, respectively.

### 2. Databases

Atomic coordinates of protein complexes are retrieved from the PDB repository^44^ and protein codes and sequences are extracted from UniProt (January 2019 release)^45^. We generate an internal database of structures with all transcription factors of the C2H2-ZF family as defined in CIS-BP database (version 1.62)^7^ and known structures in PDB. Binding information of Zinc-finger family C2H2-ZF is retrieved from bacteria one-hybrid (B1H) experiments^19^. The experiment distinguishes between Zinc-finger domains at the C-tail (F3 domain) and inner domain (F2 domain). The experiment performs the screening of all 64 possible binding sites of 3 bases characteristic of a C2H2-ZF domain. The experiment tests the interactions with multiple large protein libraries based on Zif268, with six variable amino acid positions on each individual domains F2 and F3^46^.

### 3. Interface and triads of protein-DNA structures

We define ***triads*** as a type of contacts between the protein and the double-strand DNA helix. Triads are formed by three residues: one amino acid and two contiguous nucleotides of the same strand. The distance associated with a triad is defined by the distance between the C_β_ atom of the amino acid residue and the average position of the atoms of the nitrogen-base of the two nucleotides plus their complementary pairs in the opposite strand of the helix^34^. The triad also has an associated amino acid residue number in the protein and a dinucleotide position in the DNA, defined by the sequence position of the first nucleotide of the dinucleotide (e.g. a *triad* with amino acid residue number *p*, dinucleotide in position *q* and associated distance *d* is represented as (*triad, d, p, q*)). Specific features can be added on a *triad*, defining an ***extended-triad*** *(etriad)*. These features are 1) for the amino acid: hydrophobicity, surface accessibility and secondary structure (determined with DSSP); and 2) for the dinucleotide: nitrogenous bases, the closest strand, the closest groove and the closest chemical group to the amino acid.

### 4. Statistical potentials

We use the definition of **statistical potentials** described by Feliu et al^47^ and Fornes et al.^34^ to define several **scoring functions** for the interaction between a protein and a DNA binding site using contact triads. We use the distribution at distances up to 30 Å of triads to calculate the statistical potentials. The total potential of an interaction is calculated as the sum of the potentials of all triads, or triads with their environmental features (*etriads*). In the case of *etriads*, the completeness of the reference dataset is not sufficient to sample all possible combinations. We use interactions from B1H to extend the number of interacting triads (see further details and supplementary methods). Besides, we transform the statistical potentials into Z-scores (see further), to simultaneously identify the best distance associated with a triad and the best pair formed by one amino acid and one dinucleotide.

### 5. Z-scores

The optimal condition of a statistical potential often yields a minimum. However, the minimum is not necessarily negative. The variability of signs of the potentials affects the criterion of quality of the scores. We define *Z-scores* in order to follow a criterion that incorporates the sign. We wish that the Z-score identifies simultaneously the best distance associated with a triad and the best pair formed by one amino acid and one dinucleotide. Consequently, we construct a ***zscore* function** for any type of *score* using a standard normalization with respect the average of all amino acid types (see details in supplementary).

### 6. Structural modeling of C2H2-ZF complexes

We obtain the structure of a complex by means of homology modelling using the program MODELLER^42^. The DNA binding sequence of a member of the C2H2-ZF family composed by three finger domains (F1, F2 and F3) has a length of 9 nucleotides (e.g. for Zif268). For the selection of the binding sequences associated with each finger domain we use the same sequences as in the B1H experiment^19^. The structure of Zif268 binding DNA is modelled with 23 different template structures (see details in the supplementary extension of methods). We complete the complex by modeling the structure of the DNA binding sequence. The full DNA sequence of the experiment uses 29 bases. We embed the binding site in positions 11 to 19 of this sequence. The structure of the full DNA sequence bound by Zif268 is obtained with the program X3DNA^39^ by modifying the DNA structure in the complex. We also model several structures with the complex of Zif268 binding a non-specific DNA region to be used as non-binding examples (or background). The non-binding sequence is obtained by selecting randomly a region of the sequence of the weak promoter GAL1 (see details in extended supplementary methods).

### 7. Use of experimental TF-DNA interactions to calculate statistical potentials

We use a mapping function that associates the amino acids of each hexamer sequence in the experiment of B1H with the amino acids in a template structure to derive interacting triads. Similarly, we also require a mapping between the nucleotides. For each finger (F2 and F3) and combination of 3 nucleotides, we collect all protein hexamer sequences producing significant binding signal in the B1H experiment. For the DNA sequence the mapping is on the trinucleotides of the binding-site, affecting 4 dinucleotides, while for the protein it affects 6 amino acids (see details in supplementary). Only the triads affecting the amino acids and nucleotides under test are considered for the calculation of the statistical potentials. We restrict each set of Zif268 sequences to those with highest signal of the B1H binding experiment. We define three thresholds based on the affinity percentile of a hexamer-sequence variant: 1) higher than 90%; 2) higher than 75%; and 3) higher than 50%. To calculate the affinity percentile of a variant we follow the same definition as the authors^19^. We force having around 500 DNA binding sequences per hexamer. A DNA binding sequence is repeated in proportion to the number of observations in the experiment. The contacts derived from the B1H experiment are limited to short distances (the largest contacts are around 15-20Å). Therefore, we also use contacts extracted from other structures of the C2H2-ZF family in the PDB, covering distances up to 30 Å.

### 8. Scoring TF-DNA binding

First, we calculate the interface and extract all *etriads* associated with distances shorter than 30Å. Then, the score of the interaction is defined as the sum of the scores (i.e. a specific potential) of all *etriads* with their associated distances. The same approach is applied for Z-scores. We can obtain the score of a TF without knowing the structure of the TF-DNA binary complex if it can be modelled (see details in supplementary).

### 9. Construction of PWMs using Zif268 structural models

Given the modelled structure of Zif268-DNA complex, we obtain the PWM using scores or Z-scores (as example, without loss of generality, we use the Z-score of *ES*3*DC*_*dd*_ as defined in supplementary). We collect the set of *etriads*, their associated distances, and the associated amino acid and dinucleotide positions. We obtain a **test set** with all possible DNA sequences of the binding site. We calculate the score of any sequence of the test using *ZES*3*DC*_*dd*_ (see details in supplementary for a heuristic approach when the binding size is longer than 9 bases). We normalize the scores as:

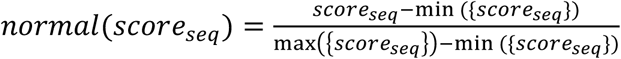

Where *score*_*seq*_ = −*ZES*3*DC*_*dd*_ and {*score*_*seq*_} is the set of all scores in the “test set”. Eventually, the normalized scores range between 0 and 1. Then, we rank the normalized scores and select only the DNA sequences producing the top scores over a **cut-off threshold** (i.e. 0.95). This produces an alignment and we use it to calculate the PWM, which we name **theoretical PWM**.

### 10. Construction of the experimental PWM

The experimental PWM of a Zif268 sequence obtained from B1H, with a specific hexamer fragment on F2 or F3, is calculated based on its affinities for different binding sites. The DNA strand of a binding site is formed by trinucleotides flanked by two fixed nucleotides (G and A for F2, and two A for F3). All binding sites targeted by a specific hexamer-fragment with affinity higher than a threshold are stored and gapless aligned to construct the PWM (e.g. the top 20% threshold uses all DNA-bound sequences with affinity percentile higher than 80%, while for a threshold of 100% we use all detected sites with any not null affinity percentile). We construct experimental PWMs for top 10%, top 25%, top 50% and for all targeted sites. These experimental PWMs are also named hexamer-specific PWMs, to distinguish from PWMs obtained with other experiments or with a different approach.

## RESULTS

### 1. Analysis of the statistical potentials

We have constructed several statistical potentials to describe the interaction between the finger domains (F2 and F3) and the DNA. We have applied a Z-score modification (see methods and further details in supplementary) on top the classical definition of potential^48^ (named PAIR). Figures 1 (A to D) show the PAIR and ZPAIR potentials between asparagine (Asn) and the dinucleotide with bases guanine-cytosine (GC) in finger domains F2 and F3. This example shows that the Z-score function preserves the optimum shortest distance, but different between domains F2 and F3. Figures 1 (E to H) show the ZPAIR potential of arginine (Arg) interacting with dinucleotides with bases adenine-guanine (AG) and cytosine-thymine (CT). Supplementary figures showing the potential PAIR and ZPAIR for all amino acids and dinucleotides can be accessed in http://sbi.upf.edu/C2H2ZF_repo. We use a set of hexamer sequence-fragments yielding affinity percentiles higher than 90%, 75% or 50% to construct the potentials. We observe that the potential is symmetric for the reversed dinucleotide (i.e. the potential resulting for the interaction of Arg with AG in Figure 1E and 1G is the same with CT in Figure 1F and 1H). However, the finger-domain has the ability to distinguish forward and reverse dinucleotides depending on structural and topological features of the DNA helix. In previous works we already developed a topological-dependent potential named ES3DC (see details in supplementary methods and in Fornes et al.^34^). The limitation of such specific potential is the completeness of the dataset, as the large number of combinations to be sampled is very high and thus requiring a large number of observations. The use of experimental data from B1H is a good opportunity to populate many triads in close distance (shorter than 20 Å) between the finger domain and the DNA binding site^19,46^. Figure 2 shows the increase of different types of contacts produced with the help of B1H data with respect to those obtained only with structures of the C2H2-ZF family in PDB^44^. Still, only some topological features of both DNA and protein conformation highlight the increase, as they are specific of the C2H2-ZF family.

**Figure 1.**
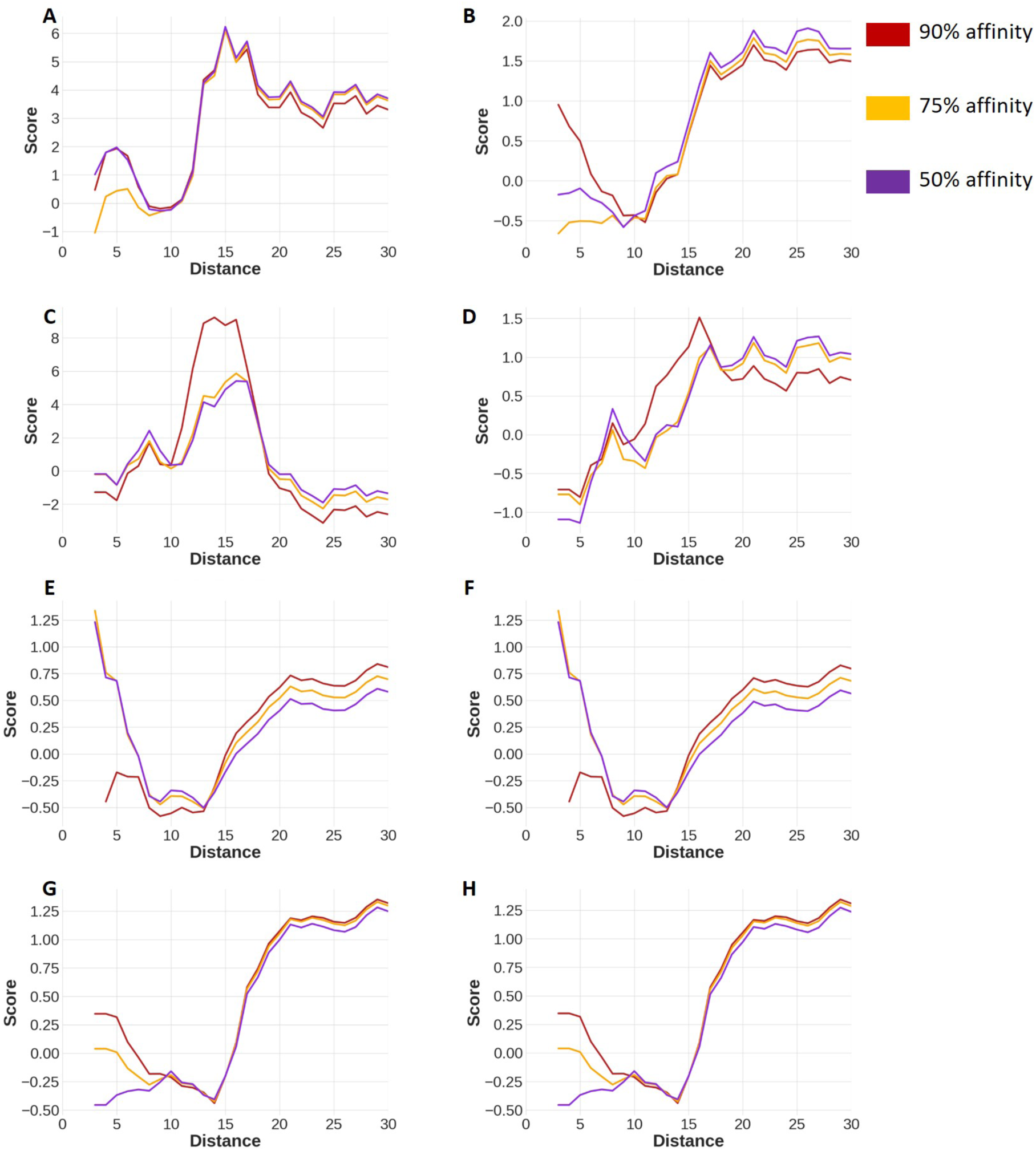
Statistical energy profiles PAIR and ZPAIR obtained with F2 and F3 domains. (**A**) Profile of Asn – GC PAIR score using F2. (**B**) Profile of Asn – GC ZPAIR score using F2. (**C**) Profile of Asn – GC PAIR score using F3. (**D**) Profile of Asn – GC ZPAIR score using F3. (**E**) Profile of Arg – AG ZPAIR using F2. (**F**) Profile of Arg – CT ZPAIR using F2. (**G**) Profile of Arg – AG ZPAIR using F3. (**H**) Profile of Arg – CT ZPAIR using F3.

**Figure 2.**
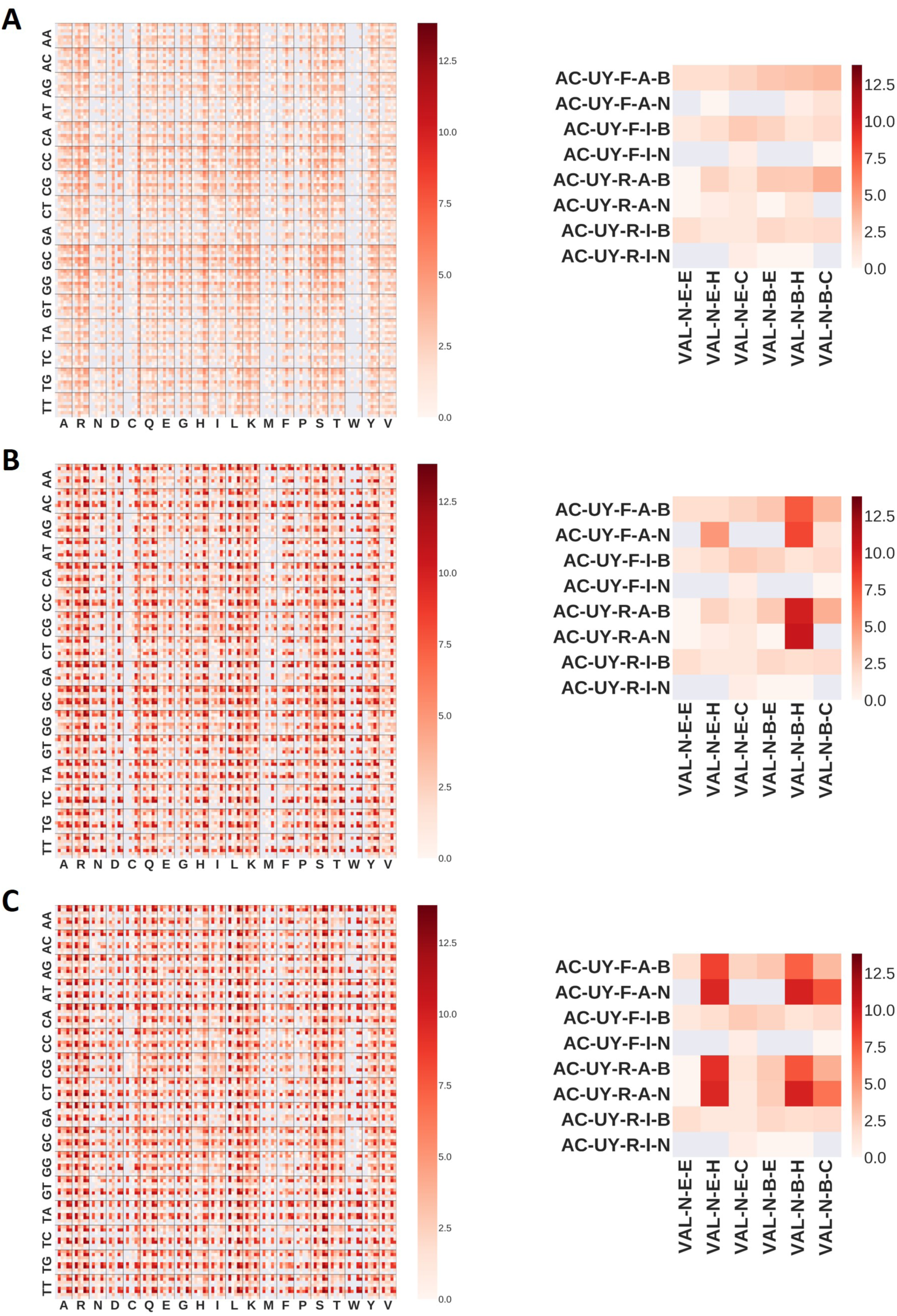
Heatmap plots of the number of amino acid – dinucleotides and their environments (etriads) at distance shorter than 30A in a logarithmic scale. Detailed view of a cell of the heatmap is shown in the right side of each heatmap. Each square inside the cell shows the number of extended-triads (in logarithmic scale) for a specific amino acid – dinucleotide (the example uses valine, Val, and adenosine-cytosine, AC) and their environments. Amino acid environments are: hydrophobicity (P as polar, N non polar), surface accessibility (E if exposed, B if buried) and secondary structure (E for β-strand, H for helix and C for coil). Dinucleotide environments are: type of nitrogenous bases (U for purine, I for pyrimidine), closest DNA strand (F for forward, R for reverse), closest DNA groove (A for major, I for minor) and closest chemical group (B if phospho-ribose backbone atoms, N if nucleobase). (**A**) Extended-triads obtained from PDB structures. (**B**) Extended-triads obtained from PDB structures and B1H experiments of the F2 domain. (**C**) Extended-triads obtained from PDB structures and B1H experiments of the F3 domain.

### 2. Prediction of PWMs in domains F2 and F3 of Zif268

To evaluate the quality of the theoretical PWMs we compare them with the results of B1H experiments^19^. We construct two types of PWMs: hexamer-specific PWMs and trinucleotide-specific PWMs. Hexamer-specific PWMs are the experimental PWMs as defined in methods, obtained by aligning all DNA trinucleotides targeted by the same amino acid hexamer. Different hexamer-specific PWMs are made for fingers F2 and F3. Trinucleotide-specific PWMs are artificial PWMs containing the combination of 3bp nucleotides (64 in total) with 100% weight in each specific position. These trinucleotides are flanked by nucleotides also specific of each assayed zinc finger (G and A for F2 and two A for F3).

We create the hexamer-specific and theoretical PWMs with the DNA binding sites of the hexamer sequences tested by B1H. We use B1H results with affinity percentiles higher than 90%, 75% and 50%. For each hexamer fragment of a finger domain, we select the binding site with highest affinity in the B1H experiment and assume that these are the three bases binding-specific of the hexamer. Theoretical PWMs are obtained by combining homology modeling and Z-scores *ZES*3*DC*_*dd*_. Structural models of the variants of Zif268 are constructed using 23 different templates (see supplementary). Hence, there are 23 theoretical PWMs for each amino acid hexamer sequence.

When comparing the theoretical PWM with the trinucleotide-specific PWMs we check the ranking position of the correct trinucleotide-specific PWM. When comparing with the experimental PWMs we use the number of nucleotide-matches to evaluate the quality. We define the number of nucleotide-matches as the number of positions where two compared PWMs share the same nucleotides with higher frequencies. Since we are focused on the trinucleotides of the binding site, we are only interested in the number of central nucleotide matches, which ranges between 0 and 3.

Using an affinity threshold of 90%, we find 131 hexamers for F2 and 82 for F3 with at least one theoretical PWM ranking the correct binding site on the top. This represents at least one hexamer sequence in 28 (for F2 domain) and 32 (for F3 domain) trinucleotide combinations out of 64. Table 1 shows the number of hexamers with three, two or one nucleotide-matches with the hexamer-specific PWM for each trinucleotide in F2 domain. Supplementary table S1 shows the results for domain F3 and additional details. Considering three or two nucleotide matches, we are able to find at least one theoretical PWM for 71% of the hexamer variants in F2 and 74% in F3. This proves that for most hexamers we are able to find a theoretical PWM with an almost perfect match with the corresponding hexamer-specific PWM. However, the selection of the template (or templates) is crucial to predict the specific binding site of a single domain. The comparisons of PWMs of all hexamer sequences tested in F2 and F3 domains with affinity percentile higher than 90% is shown in supplementary table S2.

**Table 1.**
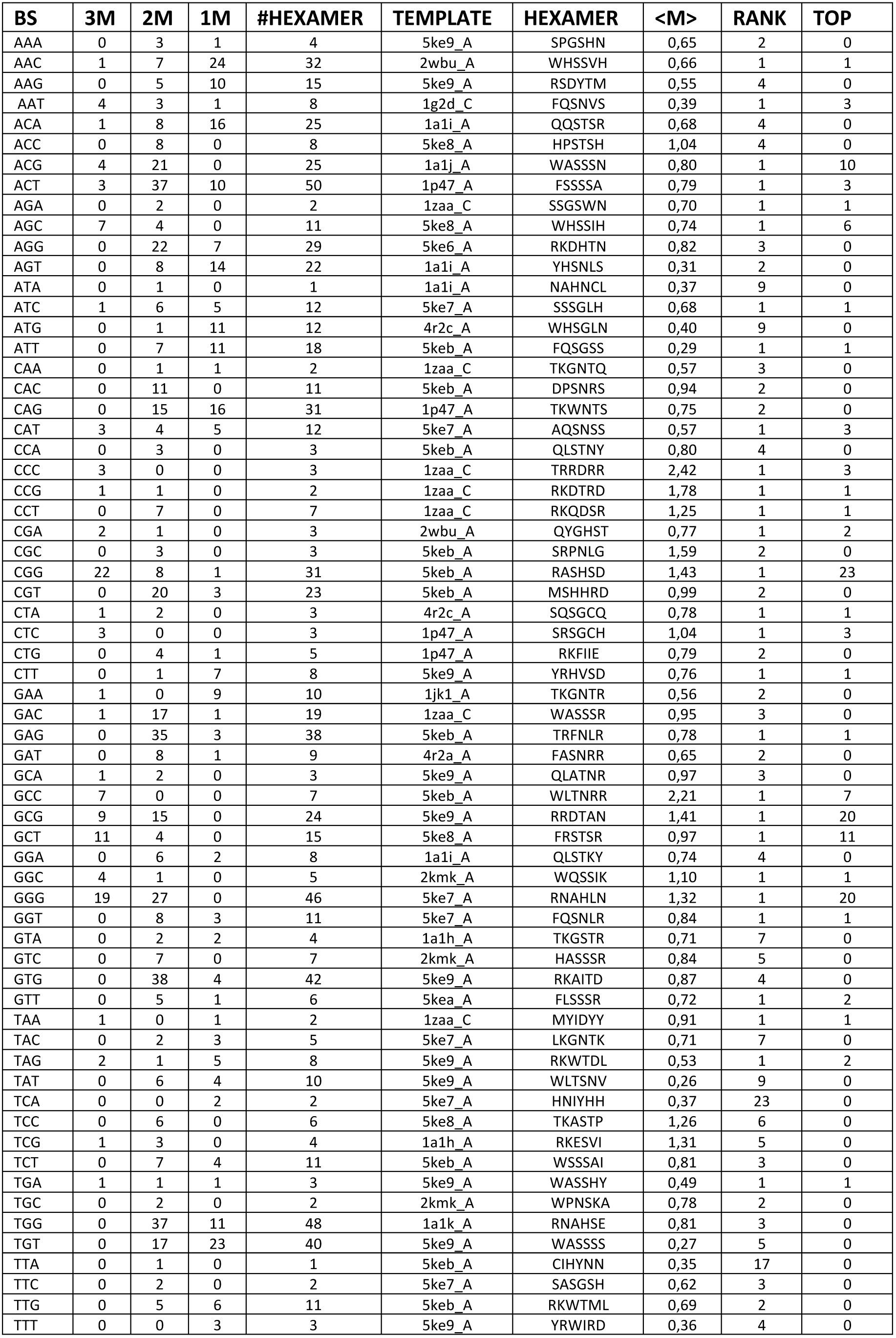
Results of the prediction of PWMs in domain F2. **BS** is the trinucleotide combination of the DNA binding site. **3M, 2M** and **1M** show the number of hexamers with at least one theoretical PWM having 3, 2 or 1 nucleotide matches with the experimental PWM, respectively. **#HEXAMER** is the total number of hexamers having as main binding the trinucleotide of the row. **HEXAMER** and **TEMPLATE** show the hexamer sequence and the code of the structure used as template for the theoretical PWM, this combination yields the highest match of nucleotides of the corresponding trinucleotide in the same row. **<M>** shows the average ratio of nucleotide matches of all theoretical PWMs and hexamer sequences with the same binding site of the row. **RANK** shows the best ranking position of the correct trinucleotide-specific PWM among all hexamers with the same binding site of the row. **TOP** shows the number of hexamers with at least one theoretical PWM ranking on the top the correct trinucleotide-specific PWM of the row.

The results obtained using affinity percentile higher than 75% and 50% are similar to use 90%. The main difference is in the number of trinucleotide combinations with successful predictions. Using an affinity threshold of 75%, around 40 trinucleotide combinations have one or more hexamers with at least one theoretical PWM ranking on top the correct trinucleotide-specific PWM, while using an affinity threshold of 50% this is around 50. Around 65-75% hexamers of F2 domain (and 75-85% for F3 domain) have 3 or 2 nucleotide-matches between the theoretical and the hexamer-specific PWM, which are similar to our previous analysis for affinity percentile higher than 90%. Details are shown in supplementary tables S1 and S2. The comparison of all PWMs are shown in http://sbi.upf.edu/C2H2ZF_repo. The fact that the quality of the theoretical PWMs is the same for all affinity percentiles analyzed suggests that our method is not able to distinguish high from low affinity binding sites; but it allows to identify approximately the TF binding site regardless of the affinity (see annex 1 in supplementary material and Figure S1).

In Figure 3 we show the comparison of some examples of hexamer-specific and theoretical PWMs grouped by zinc finger domain (all theoretical and experimental PWMs and their comparisons can be retrieved from http://sbi.upf.edu/C2H2ZF_repo). Among these examples we observe some theoretical PWMs that, although different than their expected binding sites, share common trends of the nucleotide frequencies of the experimental PWM. For example, for the binding site ATG in domain F2 by the SQSGCN hexamer (top left PWM in figure 3A), we observe similar nucleotides underlying lower frequencies between theoretical and experimental PWMs. Similarly, other examples are shown in Figure 3 with combinations of nucleotides of binding sites displaying nucleotide matches with underlying lower frequencies.

**Figure 3.**
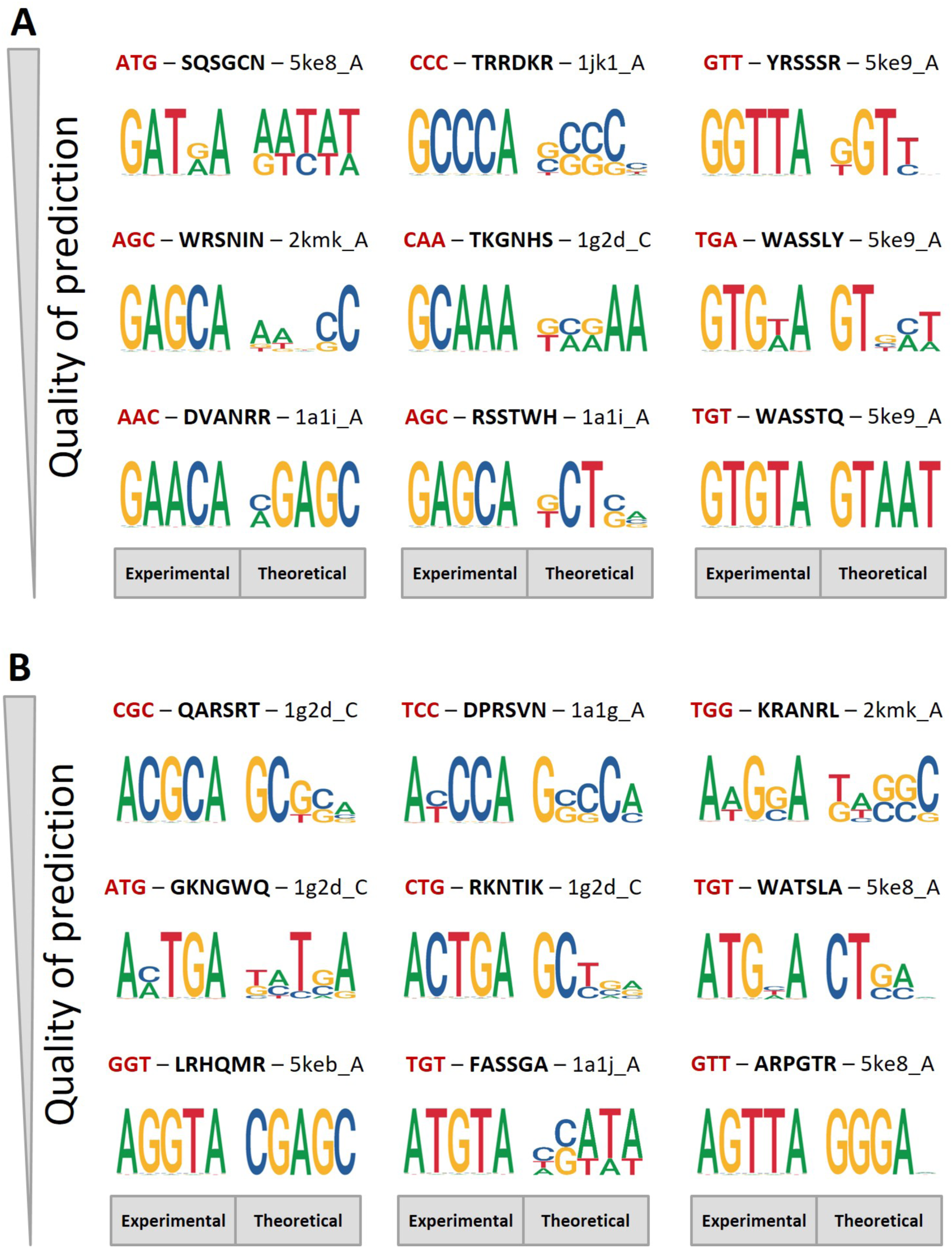
Comparison of PWMs. We compare theoretical PWMs with experimental PWMs of the same hexamer sequence variants. For each comparison we show the amino acid hexamer sequence (highlighted in bold) used to calculate the experimental PWM, the DNA binding site with highest affinity of the hexamer sequence (highlighted in red), and the PDB code of the structure used as template to obtain the theoretical PWM. (**A**) Comparison of PWMs for domain F2. (**B**) Comparison of PWMs for domain F3.

### 3. Examples of binding site prediction of C2H2-ZF transcription factors

We compare the theoretical PWMs with the PWMs retrieved from JASPAR^37^ for several TFs. We use some members of the C2H2-ZF family, composed by 3 finger-domains, with a known PWM (coded as motif) in JASPAR^37^, for which the structure of the complex with DNA is known or it can be modelled, to obtain the theoretical PWM (see supplementary table S3 and other details in supplementary material). We obtain two PWMs using statistical potentials *ZES3DC*_*dd*_ calculated with variant sequences in F2 domain (**ZES3DCF2**) and in F3 (**ZES3DCF3**) of the B1H experiment. We compare the theoretical PWMs of each TF using all contacts under 30Å, then we repeat the comparison by decreasing this threshold down to 15Å.

Figure 4 shows the JASPAR PWMs of some selected TFs, compared with the theoretical PWMs calculated with a distance threshold of 30 Å (see supplementary table S4 for more details). All theoretical PWMs and structural models can be downloaded from http://sbi.upf.edu/C2H2ZF_repo. We are able to find at least one PWM significantly similar to its motif in JASPAR (P-value <0.05 with TOMTOM) for almost all TFs (27 out of 29). The PWMs of some TFs are compared with more than one possible motif in JASPAR, often associated by some relationship in evolution (i.e. among orthologs and paralogs of different species, see details in supplementary).

**Figure 4.**
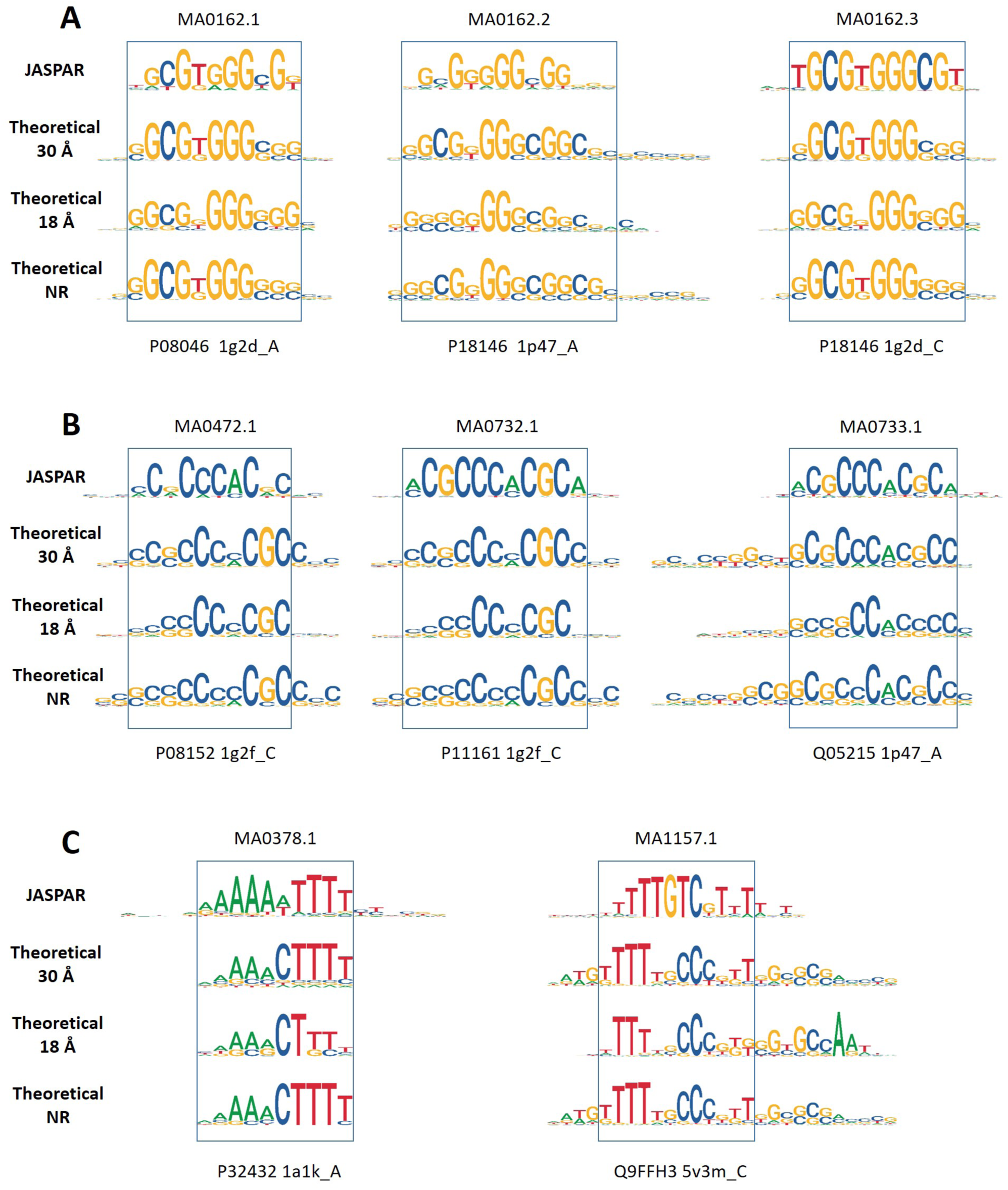
Comparison between theoretical PWMs of some members of the C2H2-ZF family and their motifs in JASPAR database. We use ZES3DCF2 statistical potentials for each TF with contacts under 30 Å, using all PDB structures of the C2H2-ZF family or avoiding those of its close homologs. A and B show examples rich on G and C nucleotides, while in C are shown examples rich on A and T nucleotides. JASPAR motifs are shown at the top of each comparison. PDB codes of the templates used to construct the theoretical PWMs are also indicated. (A) Examples of transcription factors P08046 and P18146. (B) Examples of transcription factors Q06889, P11161, Q05215 and P08152. C) Examples of P32432, Q9FFH3 and Q8H1F5.

We further test if the similarity of the TFs with the sequence of Zif268, from which the statistical potentials are derived, affects the quality of the results. We calculate the similarity as the percentage of identical residues aligned (%id) between the sequence of the TF and the sequence of Zif268. Certainly, the theoretical PWMs of TFs very similar to Zif268 are significantly similar to their motif in JASPAR. However, we also obtain theoretical PWMs for sequences with low similarity with Zif268 that significantly match with their corresponding motif in JASPAR (see supplementary). Similar conclusions are obtained when comparing the sequences of the TFs and the templates used to construct their PWMs. The bias on the statistical potential, caused by structures of close homologs to each TF query is studied in the supplementary. The main conclusion is that the bias is avoided by using contacts shorter than 18Å to construct the theoretical PWMs. We test for each TF the capacity to predict a motif in JASPAR without biases using a modification of the statistical potentials. We generate specific statistical potentials for each TF by removing the contacts of close homologs (%id >50). After avoiding the bias, we are still able to find at least one PWM significantly similar to its motif in JASPAR for almost all TFs. Also, between 11 and 14 TFs have more than 50% of the theoretical PWMs significantly similar with their motif in JASPAR, being most of them the same TFs whether close homologs are removed or not in the statistical potential (see supplementary table S4).

Not all models produce PWMs significantly similar with their corresponding motifs. For some TFs this can be explained by the low number of models produced: only one model is constructed with the length of 3-4 finger domains for Q86T24 and Q8GYC1. A detailed analysis shows that many theoretical PWMs, not significantly similar with their motif, are still able to match more than 50% nucleotide-matches with their JASPAR motif. The average ratio of identic nucleotides using all models varies between 60% and 88% for the majority of TFs (see table S4), and it is only slightly reduced after avoiding the bias.

### 4. Application to CTCF

We apply statistical potentials to predict the binding preferences of human CTCF. The DNA binding domain of human CTCF is formed by 11 zinc-finger domains of the C2H2 family (residues 266 to 577). Different DNA binding motifs have been proposed for this domain: one for the central part, flanked by one upstream and one downstream motifs^49^. The central part of human CTCF binding domain has a well-defined motif in JASPAR (MA0139.1). The structure of the complete sequence of human CTCF is unknown. However, several structures have been obtained by crystallography of different fragments bound to DNA (structures with PDB codes 5K5H, 5K5I, 5K5J, 5T00, 5T0U, 5YEL, 5YEH, 5YEF, 5UND, 5KKQ). We construct a structural model of the almost complete sequence of the binding domain of human CTCF (see supplementary figure S3). The model is constructed by superimposition of the structures 5T0U (zinc-finger domains 2-7) and 5YEL (zinc-finger domains 6-11), using the overlapping fragment of fingers 6 and 7 and removing the redundant amino acids and nucleotides from 5YEL (amino acid fragment from 455 to 512, highlighted in red in figure S3). Finger domains are shown in the protein sequence alignment and in the alignments of DNA sequences taken from the PDB structures. The C-terminal domains, taken from 5YEL, bind on the 5’ region, while the N-terminal domains from 5T0U bind on the 3’ site, thus the alignment of the DNA fragment is shown in reverse orientation with respect to the finger-domains.

Following our previous approaches, we obtain the theoretical PWM with each structure using contacts up to 30Å and the potentials ZES3DCF2 and ZES3DCF3. The PWM based on experimental data is retrieved from JASPAR, with profile motif MA0139.1, and compared with the theoretical PWMs (see details in supplementary table S5). We show in Figure 5 the logos of the JASPAR profile aligned with the logos of the theoretical PWMs. We observe a profile pattern preserved for many theoretical PWMs in which we recognize a central common region that corresponds with the JASPAR profile MA0139.1. This is the core profile of CTCF, located between nucleotides 8-22 in the forward chain (oriented form 5’-3’) of the modelled structure. Because structures such as codes 5YEL or 5UND are formed mostly by zinc-finger domains at the C-terminal region, the theoretical PWMs constructed with them match incompletely the core profile. Consequently, it is more difficult to align these PWMs, resulting in lower scores (higher P-values of TOMTOM) and small overlap. Interestingly, the profiles obtained with PDB codes 5UND, 5YEL and the 5’ site of the profile of our model, identify a pattern that has some similarities with the upstream motif mentioned by Nakahashi et al. ^49^. Motifs of flanking sites recognized by finger-domains 1-2 and 8-11 are not well-defined. Despite they are important for the recognition of the binding sites of CTCF, the current tools for motif discovery have not unraveled both downstream and upstream profiles. Consequently, we cannot test the quality of the alignments between the theoretical PWMs and many of the proposed motifs of the flanking regions, because there is no consensus.

**Figure 5.**
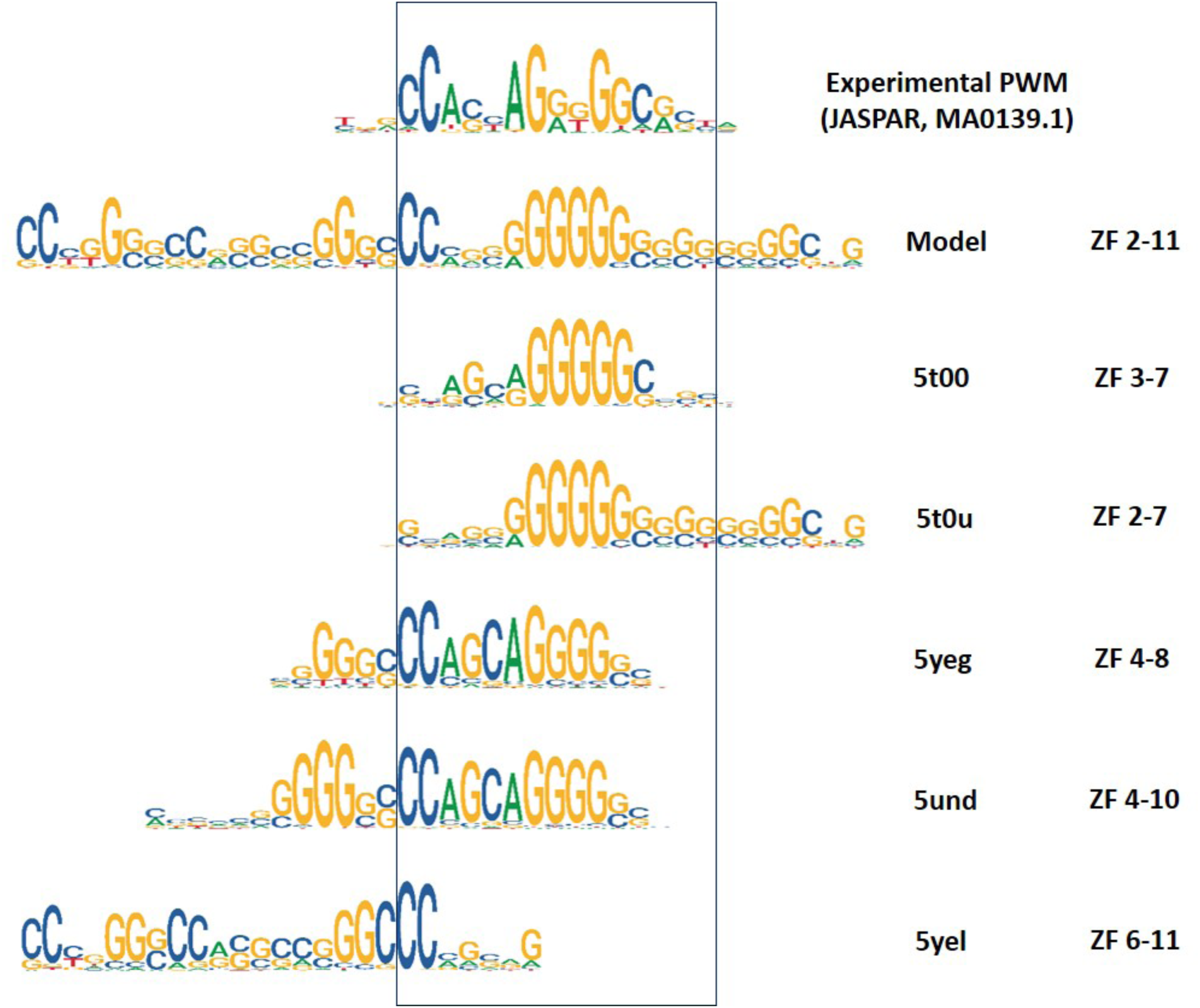
Comparison between experimental and theoretical PWMs of CTCF. The PWM on top of the figure is the experimental PWM, retrieved from the JASPAR database (MA0139.1). The rest of logos show the theoretical PWMs obtained with the model or the structures selected from PDB. The PDB code and the numbers of the zinc-finger domains corresponding to the human CTCF DNA-binding domain are shown on the right side.

## DISCUSSION

We have developed a method to predict the binding preferences of C2H2-ZF proteins using their structures, either experimentally known or modelled, to obtain one or several PWMs. Clearly, this approach is limited by the amount of structural data available. We require the structures of templates to model the proteins of the C2H2-ZF and predict the PWMs. The number of models depends on the number of templates. Consequently, the number of theoretical PWMs is larger for sequences with many templates than for those with few and this affects the capacity of the prediction.

Our analyses show that, using the scoring function *ZES*3*DC*_*dd*_ (other potentials hint towards the same conclusions), the percentage of nucleotide matches in binding sites of single-domains between theoretical and experimental PWMs is independent of the experimental affinity percentile. Therefore, although we can roughly distinguish binding from non-binding sites, we cannot distinguish intermediate degrees of affinity. This is relevant on the prediction of the effect of mutations affecting the binding strength of zinc-finger domains.

Given that our method provides several theoretical PWMs for the same TF, it entails an additional problem: selecting the correct or best PWM. However, rather than finding the best PWM from a set of theoretical PWMs, we bring the opportunity to select one among many potential solutions and help finding the binding site of a transcription factor in a DNA sequence. We proof that, for a relevant number of transcription factors that can be modelled, the number of PWMs significantly similar to an experimental PWM is larger than 50% (and the proof is valid after removing biases due to the similarity between the query sequence and the dataset used to construct the prediction). Therefore, by scanning with several theoretical PWMs of a TF, the majority of regions detected and predicted to bind will correspond with its actual binding, hitting around the right location of the binding site.

Furthermore, our approach also suggests that perhaps the same TF recognizes more than one binding site depending on its conformation. When constructing theoretical PWMs with different structures, each structure is a snapshot of the interaction between the TF and the DNA. Therefore, by using many structures we introduce the dynamic nature of proteins as an additional feature. This can be useful for C2H2-ZF proteins that may interact with DNA using different conformations or different arrangements of zinc fingers, such as CTCF. It is known that CTCF has a central binding motif plus two flanking motifs, one in downstream and another upstream^49^. CTCF binding sites display different combinations of downstream-central-upstream motifs that can be spaced by a variable number of nucleotides. Therefore, searching binding sites with many PWMs obtained from different conformations of CTCF-DNA complexes may be more informative of the whole conformational space of CTCF than a single model.

To sum up, we have developed a computational tool to predict the DNA binding preferences of C2H2-ZF proteins. With the help of homology modeling, we are able to predict PWMs for TFs for which we only know their amino acid sequence. We have tested our method by comparing theoretical PWMs with their motifs in JASPAR. We have used our approach to test PWM predictions of different regions of human CTCF and predicted a PWM to cover domains 2 to 11 of the DNA binding domain of CTCF (from downstream to upstream motifs). We offer a repository with the results and also a server to calculate the PWM using the structure of a TF as input. We think our approach may also be applied to predict the PWM of amino acid sequences of C2H2-ZF proteins that bind a specific DNA fragment. However, because of the lack of specificity on the prediction of binding affinities, further research is still needed on this goal.

## Supporting information

supplementary annex material

## ACKNOWLEDGMENTS

We thank the help of Drs. Persikov and Noyes to access and understand the data on B1H we have used in this work. This work was supported by the Spanish Ministry of Economy (MINECO) and FEDER [BIO2017-85329-R] [RYC-2015-17519]. Authors were also supported by grants: “Unidad de Excelencia María de Maeztu”, funded by the Spanish Ministry of Economy [ref: MDM-2014-0370]; IMI-JU under grants agreement no. 116030 (TransQST), resources of which are composed of financial contribution from the EU-FP7 (FP7/2007-2013) and EFPIA companies in kind contribution; The Research Programme on Biomedical Informatics (GRIB) is a member of the Spanish National Bioinformatics Institute (INB), PRB2-ISCIII and is supported by grant PT13/0001/0023, of the PE I+D+i 2013-2016, funded by ISCIII and FEDER. BO acknowledges the internationalization of the Council for the Catalan Republic. FÅ acknowledges support of Erasmus+ fellowship 2019. AM acknowledges a fellowship on Research Formation of “Generalitat de Catalunya” (FI).

