## supplementary annex material for "On the prediction of DNA-binding preferences of C2H2-ZF domains using structural models: application on human CTCF"

[1]Structural Bioinformatics Lab (GRIB-IMIM), Department of Experimental and Health Science, University Pompeu Fabra, Barcelona 08005, Catalonia, Spain, [2]Centre for Molecular Medicine and Therapeutics, BC Children's Hospital Research Institute, Department of Medical Genetics, University of British Columbia, Vancouver, BC V5Z 4H4, Canada, [3]Laboratory of Protein Design & Immunoengineering, School of Engineering, Ecole Polytechnique Federale de Lausanne, Lausanne 1015, Vaud, Switzerland, [4]Department of Biosciences, U Science Tech, Universitat de Vic-Universitat Central de Catalunya, Vic 08500, Catalonia, Spain, [5]IBERS, Institute of Biological, Environmental and Rural Science, Aberystwyth University U.K

**ANNEX OF RESULTS**

### 1. Using statistical potentials to detect binding and affinities.

We hypothesize that statistical potentials can differentiate binding from non-binding DNA sequences. To prove it, we calculate the difference of the scores obtained with the same hexamer sequence but different types of binding sites: DNA bindings sequences according to the B1H experiment, and potential non-binding sites (ten times larger in number to avoid any bias on the selection) retrieved from a background DNA fragment (see further in the extension of methods).

We use a single template to model the structure of Zif268 bound with DNA (chain A of 1P47) for this analysis. We model the sequence of DNA according to the experiment of B1H using binding and non-binding fragments (see details in methods). We use two sets of DNA sequences, one of experimentally asserted bindings and another composed of a random selection of fragments of the sequence used as background for non-binding. We restrict the scoring to the 6 variable amino acids of the finger (hexamer fragment) in interaction with the 9-mer DNA-sequence (including the 3 variable nucleotides tested for each finger sequence) at a distance shorter than 15 Å. We calculate the difference between normalized Z-scores ($ZES3DC_{dd}$) of two modelled structures obtained with two different DNA sequences but the same hexamer fragment. We define as $\Delta$score the difference between normalized Z-scores of any two models. We calculate the distribution of $\Delta$scores obtained between a bound and a non-bound DNA sequence (bound distribution). The distribution of the differences between unbound DNA sequences is used as background (i.e. this is a gaussian background distribution around zero). Supplementary figure S1 shows the distributions of $\Delta$scores for F2 and F3 domains. $\Delta$score distributions of bound and unbound are significantly different (P-value < 0.001, using a Mann-Whitney test).

### 2. Selection of TFs for the comparison with JASPAR motifs.

We identify all TFs of the C2H2-ZF family analyzed in JASPAR (a total of 181). Then, we select only those formed by 3 zinc-finger domains that can be modelled with the structure of Zif268 (even if it includes other different domains). They are identified using the HMM profile of the ZF-C2H2 family (PF00096.21) from PFAM[1], selecting a list of 40 TFs (see supplementary table S3). We use all structures of the family C2H2-ZF in the PDB[2] as potential templates and select the closest homologs to model each TF. We skip models with less than 80 amino acids and only analyze proteins with 3-4 finger domains (i.e. less than 120Aa), which reduces the set to 29 TFs. No more than 40 templates are used per TF, usually between 10 and 30 and occasionally only one or two, so every TF has several models (one per template). We use the models to obtain the theoretical PWMs of each TF using all contacts under 30Å (see details in methods). For each TF we compare its motif in JASPAR with the set of theoretical PWMs. We also test the theoretical PWMs with contacts under different thresholds (ranging between 15Å to 30Å).

### 3. Examples of the prediction of PWMs among ortholog and paralog TFs.

The PWMs of some TFs are compared with more than one possible motif in JASPAR, often associated by some relationship in evolution (i.e. among orthologs and paralogs of different species). Examples are shown in figure S2. For example, Q43474 and Q60793 have PWMs created with templates 1p47 (chain C), 1a1i (chain A) and 1a1j (chain A) respectively similar with motifs MA0039.2, MA0039.3 and MA0039.1; or some models of P08046 and P18146 generate PWMs very similar to motifs MA0162.1, MA0162.2 and MA0162.3. Actually, P18146 corresponds to the human sequence of the protein produced by gene EGR1, P08046 corresponds to the sequence of its ortholog gene in mouse, while motif MA0162.1 corresponds to the mouse gene and MA0162.3 and MA0162.2 are the motifs in JASPAR corresponding to human. The high similarity of all of them is caused by the similarity of the DNA binding sequence, which is reflected in the structural models. Another example is shown in figure S2 to compare different motifs (MA0472.1, MA0732.1 and MA0733.1) with human and mouse sequences of non-orthologous genes EGR3 and EGR2 (Q06889 and P08152), respectively.

### 4. Study of biases caused by the sequence similarity between the transcription factor and the template when comparing a theoretical PWM with its motif in JASPAR.

The statistical potentials are obtained with structures of C2H2-ZF members and experiments from B1H data of C2H2-ZF proteins. Here we test if the success of our approach depends on the similarity between the query sequence and the sequences of the structures used to create the PWMs and the statistical potentials. First, we compare the sequence of the query with the sequence of Zif268, because this has been used in all the models to create the potentials using B1H experiments. Second, we compare the sequence of the query with the sequence of its template (closest homolog with available structure). These templates are used to generate the statistical potentials as well as to make the models from which we will obtain the theoretical PWMs. We calculate the similarity between the theoretical PWM and its motif in JASPAR. We measure this similarity as the logarithm of the P-value of significance. Then, we compare the sequences of the query and the template and calculate their similarity as the percentage of identical residues in the alignment. Finally, we compare the criteria of similarity to find if there is a relationship between both.

Figure S3, plots A and B, show the comparison of sequence and PWMs similarities with respect to the sequence of Zif268. We observe a strong Pearson correlation statistically significant ($P < 0.001$) using ZES3DCF2 (0.725) and ZES3DCF3 (0.74) statistical potentials. This correlation is mainly caused by theoretical PWMs very similar to their motif in JASPAR obtained when the sequences are almost identical to Zif268, while the theoretical PWM deviates from the corresponding motif for proteins with low sequence similarity with Zif268. After removing proteins highly similar to Zif268 (> 60% sequence identity) the Pearson correlation is downgraded. Similarly, plots C and D in Figure S3, show the sequence and PWMs similarities where sequence similarities are calculated between the sequence of the TF and the template used to construct the theoretical PWM. Strong and significant Pearson correlations are also observed using ZES3DCF2 (0.70) and ZES3DCF3 (0.68) statistical potentials. Nevertheless, we also observe that TFs with sequence very different to the sequence of Zif268 yield theoretical PWMs very accurate (significantly similar to the corresponding motif in JASPAR). Similarly, theoretical PWMs of TFs, constructed with templates very different in sequence, can be very similar to their motif in JASPAR. These results demonstrate the applicability of the approach using the structures of remote homologs, or templates with low sequence similarity.

Further, we have tested the bias effect on the statistical potentials for the application on human CTCF. The number of structures of CTCF used to calculate the statistical potentials is about 12% of the total of C2H2-ZF structures (13 out of 113). These structures mostly affect amino acid-dinucleotide interactions at distances between 15Å and 30Å. This may have a significant effect on the potential, so we have removed these structures and recalculated the statistical potentials, which are then free of any information on CTCFs.

We compare the theoretical PWMs of the modelled structure of CTCF with motif MA0139.1. This structure contains zinc fingers 2 to 11 and its corresponding PWMs are obtained with statistical potentials ZES3DCF2 (and ZES3DCF3) at cut-off thresholds between 15 and 30 A. Besides, these PWMs are made using two types of potentials: potentials including all C2H2-ZF structures and potentials that do no include CTCF structures.

The best matches with the JASPAR motif of CTCF (MA0139.1) are obtained with cut-off distances between 15Å and 22Å instead of 30Å when the potential doesn't includes structures of CTCF. Figure S5 shows the comparison of logos. We observe that the inclusion of CTCF structures is only noticed after 20Å, while the PWMs between 15Å and 20Å are mostly influenced by the B1H experiments, showing that the theoretical PWMs calculated with ZES3DCF2_all or ZES3DCF2_CTCFfree are more similar between them, but less with respect to motif MA0139.1. This test suggests that we can avoid the biases on the statistical potential, caused by structures of close homologs of a testing TF, by reducing the contacts to less than 20Å. Further, we quantify this effect by comparing the ratio on nucleotide-matches between the JASPAR motif and each theoretical PWM with respect to the cut-off distance used to calculate contacts (see table S5).

Theoretical PWMs of TFs used in the comparative analysis with motifs in JASPAR (section 3 of the manuscript) are obtained with potentials ZES3DCF2 and ZES3DCF3 using cut-off distances at 15 Å, 18 Å, 22 Å, 25 Å, 28 Å and 30 Å and can be downloaded from http://sbi.upf.edu/C2H2ZF_repo. Supplementary table S4 shows the results using contacts under 18 Å. These results prove that for 8 (using ZES3DCF2) and 13 (using ZES3DCF3), out of 29 TFs, the number of theoretical PWMs significantly similar to the corresponding motif is higher than 50%. These are very significant results, because it suggests we may find the potential binding of a TF while reducing the biases on the statistical potential with a relevant degree of accuracy. However, the number of TFs is lower than when we use statistical potentials with all contacts under 30 Å. Therefore, we have tested the approach using all contacts, but with unbiased statistical potentials.

To obtain unbiased statistical potentials, we first identify for all TFs under test the structures in the database of their close homologs. Then, we produce TF-specific statistical potentials without the contacts derived from their homologs in the database. For example, TFs P08046, P18146, Q06889, P08152, P11161, Q05215 are too similar (more than 50% identical residues aligned) with the sequences of the known structures of Zif268 (i.e. PDB codes 1A1J, 1AAY, 1ALI, 4X9J, 1A1K, 4R2A, 1ZAA, 1G2F, 1A1F, 1G2D, 1A1L, 1A1G, 4R2C, 1A1H). Consequently, we construct a specific statistical potential without using contacts from these structures (except those obtained by means of B1H experiments). The results are shown in tabs 5 and 6 of table S4. They proof a good consistency with the predicted PWMs when we use all known structures, with only slight decrease on the ratio of aligned nucleotides and in the number of theoretical PWMs that are significantly similar with their corresponding motif in JASPAR.

### LEGENDS FOR SUPPLEMENTARY FIGURES

**Figure S1**. Density plot (ratio) of the difference between scores of binding and non-binding interactions ($\Delta$score). Scores are calculated with hexamers in F2 (A) and F3 (B) using their binding sites and artificially constructed non-binding sites. Distribution of the differences between binding and non-binding interactions are shown in red, differences between non-binding interactions are shown in grey.

**Figure S2.** Comparison between some theoretical PWMs of C2H2-ZF and their motifs in JASPAR database for ortholog and paralog examples. We use statistical potentials ZES3DCF2 to generate the theoretical PWMs. JASPAR motifs are shown at the top of each comparison. PDB codes of the templates used to construct the theoretical PWMs are also indicated. We highlight with red boxes the theoretical PWMs corresponding to the actual motif of each sequence associated with the same species (only for examples of similar motifs). (A) Examples of transcription factors with more than one motif in JASPAR for orthologous genes (P08046 corresponds with the mouse associated motif MA0162.1 and P18146 corresponds with human associated motifs MA0162.2 and MA0162.3). (B) Examples of transcription factors with different, but very similar motifs in JASPAR, using non-orthologous genes, for proteins Q06889, P11161, Q05215 and P08152 (P08152 corresponds with the same species of MA0472.1).

**Figure S3.** Scatter plots for the comparison of sequence and PWM similarities. The similarity between the theoretical PWM of a TF and its JASPAR motif is used to score the quality of the prediction. This is calculated as -$Log_{10}$(P-value), where P-value is obtained with TOMTOM from the comparison of both PWMs. The similarity of the sequence of a TF and the template used to construct the theoretical PWM (or the sequence of Zif268) is calculated as the percentage of identical residues in the alignment. Points in the scatters show the quality of the prediction (i.e. similarity between PWMs) versus the similarity of the sequences. In A and B, we compare the quality of the prediction of the PWM of a TF versus the similarity of its sequence with Zif268, where the theoretical PWM is calculated with the statistical potential ZES3DCF2 (A) or ZES3DCF3 (B). In C and D, we compare the quality of the prediction of the PWM of a TF versus the similarity between the sequence of the TF and the template used to construct the theoretical PWM, with statistical potentials ZES3DCF2 (C) and ZES3DCF3 (D). Lines in red show the linear square fitting of the Pearson correlation between sequence and PWM similarities. Lines in blue show the same after removing the points obtained for sequence similarities larger than 60%. For each fitting line we show the coefficient of the Pearson correlation (r) and its corresponding P-value (p).

**Figure S4**. Details of the modeling of human CTCF DNA-binding domain. (**A**) Alignment between the sequence of human CTCF DNA-binding domain, the sequences of selected PDB structures of CTCF DNA binding domain and the sequence we use to construct the model. The model of human CTCF DNA-binding domain is constructed with the superposition of the structures of 5T0U and 5YEL. The fragments of the sequences of 5T0U and 5YEL used for the superposition are highlighted in red. Zinc finger domains 2 to 11 are highlighted in the alignment: Zinc-finger domains of the sequence model aligned with the sequence of the structure 5T0U are shown in red, zinc-finger domains aligned with the sequence of the structure 5YEL are shown in blue and zinc-finger domains aligned with both are shown in purple. (**B**) Alignment between the DNA sequences of the PDB structures and the sequence of DNA which structure is modelled. Nucleotides of the binding site of each finger domain are highlighted in boxes and numbered after each binding domain. (**C**) Ribbon plot of the modelled structure of the human CTCF-DNA complex showing zinc fingers 2 to 11.

**Figure S5**. Comparison of theoretical PWMs obtained with potentials ZES3DCF2_all (using all known structures of C2H2ZF proteins, including structures of CTCF) and ZES3DCF2_CTCFfree (free of CTCF structures). Theoretical PWMs are obtained at cut-off thresholds of 15Å, 17 Å, 19Å, 22Å, 24Å, 28Å and 30Å. Motif MA0139.1 of CTCF is aligned with the theoretical PWMs.

**LEGENDS FOR SUPPLEMENTARY TABLES**

**Table S1.** Comparison of hexamer-specific and theoretical PWMs grouped by DNA binding sites. Each tab corresponds to a threshold of percentage of affinity (90%, 75% or 50%), used to obtain the results. All tabs have the data organized in the same columns.

**Column A** shows the zinc finger used to get the potentials (either F2 or F3). **Column B** shows the DNA binding site. **Columns R to U** show the number of hexamers with a maximum number of nucleotide matches with the experimental hexamer-specific PWM (nucleotide matches can be 3, 2, 1 or 0; respectively for each column). **Column X** shows the total number of amino acid hexamers tested to bind the corresponding DNA binding site. **Columns Y to AA** show the percentage of hexamers with a maximum number of nucleotide matches with the experimental hexamer-specific PWM (nucleotide matches can be 3, 2 or 1; respectively for each column). **Column AB** shows the hexamer sequence for which we find a theoretical PWM with the highest number of nucleotide matches with the experimental hexamer-specific PWM. **Column AC** shows the structural model that produces the theoretical PWM in column AB, identified by the PDB code of the template, the hexamer sequence, and the finger domain. **Column AF** shows the average of nucleotide matches between all theoretical PWMs and all hexamer-specific PWMs with the same binding site of the row. **Column AG** shows the average score for all theoretical PWMs with the same binding site of the row. This score is obtained as the minus logarithm of the P-value obtained by TOMTOM in the comparison between theoretical and experimental hexamer-specific PWMs. **Column AI** shows the minimum ranking of the trinucleotide-specific PWM using all theoretical PWMs of hexamer sequences with the same binding site of the row. **Column AJ** shows the average ranking position of the trinucleotide-specific PWM using all theoretical PWMs of hexamer sequences with the same binding site of the row. **Column AK** shows the number of theoretical PWMs with the same binding site of the row that rank the trinucleotide-specific PWM in the top. **Column AL** shows the percentage of theoretical PWMs that are significantly similar to their corresponding experimental hexamer-specific PWM according to TOMTOM (P-value < 0.05). **Column AM** is one if at least one theoretical PWM among all hexamer-specific PWMs with the same binding site of the row ranks in the top the trinucleotide-specific PWM, and zero otherwise. **Column AN** is one if at least one theoretical PWM matches three nucleotides with the DNA binding site, and zero otherwise. **Column AO** is one if at least one theoretical PWM matches two nucleotides with the DNA binding site, and zero otherwise. **Column AP** is one if at least one theoretical PWM matches three or two nucleotides with the DNA binding site, and zero otherwise.

**Table S2.** Comparison of hexamer-specific and theoretical PWMs. Each tab corresponds with a threshold of percentage of affinity (90%, 75% or 50%) used to obtain the results. All tabs have the data organized in the same columns. **Column A** shows the hexamer sequence. **Column B** shows the zinc finger domain where the hexamer is embedded (either F2 or F3). **Column C** shows the trinucleotide of the binding site. **Column D** shows the enlarged DNA binding site by the flanking nucleotides. **Column P** shows the PDB template of the model that produces the PWM ranking the trinucleotide-specific PWM in the minimum position. **Column Q** shows the minimum position of the ranking of the trinucleotide-specific PWM. **Column T** shows the average ranking position of the trinucleotide-specific PWM calculated with all theoretical PWMs of the same hexamer models. **Column W** shows the model that produces the theoretical PWM that has the maximum number of nucleotide matches with the hexamer-specific PWM. **Column X** shows the maximum number of nucleotide matches between the theoretical PWM in W and the experimental hexamer-specific PWM. **Column Y** shows the average of nucleotide matches between the 23 theoretical PWMs constructed with the models and the experimental hexamer-specific PWM. **Column AB** shows the model that produces the theoretical PWM with maximum score when comparing it with the experimental hexamer-specific PWM. **Column AC** shows the maximum score obtained with the TOMTOM comparison between the theoretical PWMs and the experimental hexamer-specific PWM. **Column AD** shows the average of the scores obtained with the TOMTOM comparison of the theoretical PWMs and the experimental hexamer-specific PWM**.** **Column AE** shows the lowest P-value from the comparisons between the theoretical PWMs and the experimental hexamer-specific PWM.

**Table S3.** Selection of C2H2-ZF proteins from the JASPAR database. Tab 1 shows the JASPAR ID and the uniprot ID for all proteins from the JASPAR database that belong to the C2H2-ZF family. Tab 2 and tab 3 show the results of the selection of proteins from the JASPAR database that according to a PFAM search have zinc finger domains. In tab 2 all proteins with at least one zinc finger domain are shown. In tab 3 results for selected proteins with only 3-4 zinc finger domains. In tab 2 and tab 3 data is organized with the same columns. **Column A** shows the E-value of the match between the PFAM model and the whole sequence. **Column B** shows the score of the match for the whole sequence. **Column C** shows a correction term applied to the score depending on the bias on sequence composition on the overall sequence. **Column D** shows the E-value of the best match between PFAM C2H2-ZF domain and the best matched region of the sequence. **Column E** shows the score of the best match between PFAM C2H2-ZF domain and the best matched region of the sequence. **Column F** shows a correction term applied to the score depending on the bias on sequence composition for the sequence region having the best match with the HMM. **Column G** shows the number of C2H2-ZF domains found with the PFAM model. **Column H** shows the natural number of column G. **Column I** shows the uniprot ID of the sequence studied.

**Table S4.** Comparison of theoretical PWMs of transcription factors with their motifs in the JASPAR database. Tab 1 shows results obtained using statistical potentials based on data for domain F2 (ZES3DCF2) and cut-off distance of 30 Å. Tab 2 shows results obtained using statistical potentials based on data for domain F3 (ZES3DCF3) and cut-off distance of 30 Å. Successive odd tabs show results with statistical potential ZES3DCF2, while even tabs show results with statistical potential ZES3DCF3. Tabs 3 and 4 are obtained using the corresponding potentials and cut-off distance of 18 Å, tabs 5 and 6 are obtained with TF-specific statistical potentials where contacts from close homologs have been removed. In all tabs the information is structured in the same columns. **Column A** shows the UniProt ID of the protein. **Column B** shows the JASPAR motif ID. **Column D** shows the model that produces the PWM with the largest number of nucleotide matches with the PWM of JASPAR. The code identifies the structural model by indicating: the UniProt ID, the first and last residue of the modelled structure and the PDB code of the template (i.e. P08046:333:420_1g2d_C_1 is the model of P08046, between amino acids 333 and 420, constructed with chain C of 1g2d as template). **Column E** shows the P-value of the theoretical PWM selected in D and the JASPAR PWM. **Column K** shows the number of models that produce a theoretical PWM significantly similar to JASPAR PWM (P-value < 0.05). **Column L** shows the number of complete models for the DNA binding domain of the protein under analysis. **Column M** shows the total number of models (complete or incomplete) for the DNA binding domain of the protein under analysis. **Column N** shows the average of the minus logarithm of the P-values of all theoretical compared with JASPAR motif. **Column O** shows the average of the number of nucleotides that match the JASPAR PWM. **Column P** shows the ratio of nucleotide matches over all nucleotides in the binding motif of JASPAR for all comparisons with theoretical PWMs. **Column Q** shows the ratio of models that produce theoretical PWMs that are significantly similar to the JASPAR PWM (P-value < 0.05). **Column S** is one if the ratio in Q is higher than 0.5 and a zero otherwise. Rows showing redundant motifs of sequences already analyzed on top are highlighted in yellow.

**Table S5.** Comparison between theoretical PWMs of CTCF and MA0139.1 motif in JASPAR. Tab 1 shows results obtained using statistical potentials that include known the structures of CTCF in PDB. Tab 2 shows results obtained using statistical without contacts from structures of CTCF. In both tabs the information is structured in the same columns. **Column A** shows the JASPAR motif. **Column B** shows the model, potential and maximum distance to calculate the contacts and create the theoretical. An example of the code is 5und_F2_15, for a theoretical PWM obtained with the structure of 5UND from PDB, using potential ZES3DCF2 and cut-off distance of 15 Angstroms. **Column C** shows the orientation of the match between the theoretical and the experimental PWMs. If it is "+" it means that both PWMs are on the same orientation (i.e. both forward), if it is "-" it means that one of the PWMs must be reversed (i.e. one forward and the other reverse). **Column D** shows the TOMTOM P-value of the comparison between the theoretical PWM and the JASPAR motif. **Column E** shows the score of the comparison between the theoretical PWM and the JASPAR motif. **Column F** shows the offset between the theoretical PWM and the JASPAR motif once aligned. **Column G** shows the number of aligned positions between the theoretical PWM and the JASPAR motif. **Column H** shows the number of nucleotide matches between the theoretical PWM and the JASPAR motif. **Column I** shows the ratio of nucleotide matches between the theoretical PWM and the JASPAR motif with respect to the shortest binding site. **Column J** shows the consensus DNA sequence for the JASPAR motif. **Column K** shows the consensus DNA sequence for the theoretical PWM.

### EXTENSION OF METHODS

#### 1. Software requirements

We require the following software: DSSP (version CMBI 2006) [3] provides protein structural features; X3DNA (version 2.0) [4] is used to analyze and generate DNA structures; *matcher* and *needle*, from the EMBOSS package (version 6.5.0) [5], produces local and global alignments, respectively; BLAST (version 2.2.22) [6] is employed to search homologs of a target protein; MODELLER (version 9.9) [7] is used to create structural models with all the templates similar to our target; and the programs FIMO and TOMTOM from the MEME suite[8] are used to scan a DNA sequence with a Position-Weight Matrix and to compare two PWMs, respectively.

### 2. Databases

Structural information is retrieved from the PDB repository [2] and protein codes and sequences are extracted from UniProt (January 2019 release) [9]. We select all transcription factors of the C2H2-ZF family as defined in CIS-BP database (version 1.62) [10] with known structures in PDB to generate the internal database of structures (set **PDB$_{\mathbf{DNA}}$**). We rearrange the set of structures by separating them in chains and constructing a set of structures formed by single protein-chains interacting with a double-strand helix (**single-chain PDB$_{\mathbf{DNA}}$**).

Binding information of Zinc-finger family C2H2-ZF is retrieved from bacteria one-hybrid (B1H) experiments [11]. The experiment distinguishes between Zinc-finger individual domains at the C-tail (F3 domain) and inner domain (F2 domain). The experiment performs the screening of all 64 possible 3bp targets for interactions with C2H2-ZF domains from multiple large protein libraries based on Zif268 structure with six variable amino acid positions on each individual domains F2 and F3 [12].

### 3. Interface and triads of protein-DNA structures

The interface between a transcription factor and DNA is defined by the residues (amino-acids and nucleotides) in contact. A general approach for protein-protein interactions is to consider that two residues are in contact if the distance between a pair of atoms from each residue is shorter than 5Å. We define ***triads*** as a type of contacts between the protein and the double-strand DNA helix. Triads are formed by three residues: one amino-acid and two contiguous nucleotides of the same strand. The distance associated with a triad is defined by the distance between the $C_\beta$ atom of the amino acid residue and the average position of the atoms of the nitrogen-base of the two nucleotides plus their complementary pairs in the opposite strand of the helix [13]. The triad also has an associated amino-acid residue number in the protein and a dinucleotide position in the DNA, defined by the sequence position of the first nucleotide of the dinucleotide.

We define the **interface** between a transcription factor and DNA as: the set of *triads* with associated distances shorter than 15 Å, and their associated amino-acid residue number and dinucleotide position (e.g. a $triad$ with amino-acid residue number $p$, dinucleotide in position $q$ and associated distance $d$ is represented as $(triad, d, p, q)$).

Specific features can be added on a *triad*, defining an ***extended-triad***:

1) Hydrophobicity of the amino-acid. Amino-acid residues are split in **Polar (P):** {Arg, His, Lys, Asp, Glu, Ser, Thr, Asn, Gln, Cys, Gly} and **Non-polar (N):** {Ala, Ile, Leu, Met, Val, Phe, Trp, Tyr, Pro}.
2) Surface accessibility of the amino-acid. We use DSSP to calculate the percentage of accessibility of the residue in the unbound structure of the protein. If the percentage is smaller than 50% the amino-acid is **buried (B)**, otherwise it is **exposed (E)**.
3) Secondary structure of the amino-acid. We use DSSP to calculate the secondary structure of the protein. The amino-acid of the triad is either in regular secondary

structure (**H** if in **α-helix**, **E** if in **β-strand**), or in a **non-regular** secondary structure (**C**)

4) Nitrogenous bases: We classify nucleotides by their nitrogenous bases in two types, **purines (U):** {A, G} and **pyrimidines (Y):** {C, T}.
5) Closest strand. We use X3DNA to define the strands **forward** and **reverse** of the DNA. Next, we calculate the distance of all atoms of the two nucleotides to the $C_\beta$ of the amino-acid. We define the strand closest to the amino-acid (i.e. with the atom at minimum distance) as either the strand of the two nucleotides of the triad or the strand of their complementary pair in the opposite strand, which can be either **forward (F)** or **reverse (R)**.
6) Closest Groove. We calculate the distances between the $C_\beta$ of the amino-acid and the closest phosphates of the dinucleotides in both strands (i.e. the strand of the two nucleotides of the triad and its complementary). We calculate the positions of the closest phosphates in both strands (let be $P_f$ and $P_r$, backbone phosphates of nucleotides f and r, respectively). We select the closest phosphate of both and its corresponding strand. Let assume that $P_f$ is the closest phosphate and define its strand as "s", being "S" the opposite strand. Then, we consider the set of backbone phosphates in "S" around the position complementary of nucleotide "f" (6 nucleotides up and down). Depending on their distance to "f" (towards 22Å is a major groove and towards 12 Å a minor groove), we classify them as part of the minor or major groove with respect to nucleotide "f". This is a classification of 12 nucleotides around the complementary of "f" in two groups: 1) set at large distance (i.e. major groove); and 2) set at short distance (i.e. minor groove). Necessarily, $P_r$ is in the list classified in **major** or **minor groove**. We use the classification of $P_r$ to define the type of the closest groove of the amino-acid (i.e. we should say that this is the groove faced by the amino-acid, defined by the pair $P_f$ and $P_r$, in closest proximity to the amino-acid). The closest groove is defined as **major groove (A)** if $P_r$ is in the list classified in major, otherwise it is defined as **minor groove (I).**
7) Chemical group of the nucleotides. We distinguish two main chemical groups of each nucleotide, the nitrogenous base (**N**) and the backbone (**B**) that includes the phosphate and sugar. We calculate the distances between the $C_\beta$ of the amino-acid and the atoms of the two nucleotides and their complementary. We select the atom with the shortest distance as the closest atom between the nucleotides and the amino-acid. We define the chemical group of the nucleotides of the triad as the chemical group to which belongs the closest atom (i.e. **N** or **B**)

Added features of triads can also be used on their own as **feature-triads** (or environment triads), and every extended-triad has an associated feature-triad, both associated with the same distance, amino-acid number and dinucleotide position. As an example, let be a lysine residue and two nucleotides, adenosine and guanosine, forming the triad [K,(AG)] at 15.6Å, with lysine in residue number 32 and adenosine in 5, described as ([K,(AG)], 15.6, 32, 5). If lysine surface is mainly exposed to solvent, in a α–helix conformation and the closest strand of DNA is the forward strand, the closest atom of the two nucleotides is a phosphate and the amino-acid faces the minor groove, the extended-triad is [{K,(p-H-E)},{(AG),(UU-F-I-B)}], where added features are (p-H-E) for the amino-acid and (UU-F-I-B) for the dinucleotide. This produces a feature-triad defined as [(p-H-E),( UU-F-I-B)] at 15.6Å.

We require to define some functions on the sets of triads, extended-triads and feature-triads to extract some of the values collected from a complex structure and apply other functions:

$$f_{td}(triad, d, p, q) = (triad, d)$$
$$f_t(triad, d, p, q) = (triad)$$
$$f_d(triad, d, p, q) = (d)$$
$$f_a(triad, d, p, q) = (p)$$
$$f_n(triad, d, p, q) = (q)$$

The same functions are applied to extended-triads and feature-triads accordingly modified. We also define functions to substitute some of the elements of a triad, extended-triad or feature-triad (the example is given for *etriads* without loss of generality):

1) $\varepsilon_a(etriad, r)$ is a function that substitutes amino-acid residue "a" of the *etriad* by amino-acid-residue "r", with the corresponding change of hydrophobicity but preserving the rest of features and measures associated with the triad.
2) $\eta_v(etriad, n)$ is a function that substitutes dinucleotide $v$ by $n$, in $\Lambda = \{A, C, G, T\} \times \{A, C, G, T\}$, with the corresponding change of nitrogenous bases and preserving the rest of features and measures associated with the triad.

### 4. Statistical potentials

We use the definition of **statistical potentials** described by Feliu et al [14] and Fornes et al. [13] to define several **scoring functions** for the interaction between a protein and a DNA binding site using contact triads. We use triads, their associated measures (i.e. distance, amino-acid number and dinucleotide position) and their added features to calculate the frequencies per distance in bins of 1Å (i.e. intervals [0,1], [1,2], [2,3], [3,4], [4,5] etc.) up to 30 Å. We also calculate the frequencies using the distance as cut-off (i.e. the frequency of triads at distance shorter than "d", with d= 1,2,3,4, etc.). To obtain the frequencies, we first calculate the size (cardinality, defined by the function "Card") of the sets of triads, associated with a distance (d), taken from the set of structures of protein-DNA interactions ($PDB_{DNA}$) and grouped by their associated distance, limited to a maximum of 30Å (i.e. with $i \in [1,30]$). The set of triads, associated with distances, amino-acid residue-number and dinucleotide position, is named ***3Dset***. Then, frequencies are defined using functions defined in section 3 as follows:

$N_D(triad, i) = Card(\{f_{td}(x) | where\ x \in 3Dset\ and\ (i-1) < d \leq (i)\})$ (eq. 1)
$N_c(triad, i) = Card(\{f_{td}(x) | where\ x \in 3Dset\ and\ 0 < d \leq i\})$ (eq. 2)

Where $x = (triad, d, p, q)$ is a triad associated with a distance d, amino-acid residue-number $p$ and dinucleotide position $q$, taken from the set *3Dset*, $N_D$ is defined using bins and $N_c$ using cut-offs. Similarly to ***3Dset*** we define the sets ***e3Dset*** and ***f3Dset*** for extended-triads (*etriad*) and feature-triads (*ftriad*), and calculate $L_D$, $L_c$, $M_D$ and $M_c$ with $i \in [1,30]$ as:

$L_D(etriad, i) = Card(\{f_{td}(x)|where\ x \in e3Dset\ and\ (i-1) < d \leq (i)\})$ (eq. 3)
$L_c(etriad, i) = Card(\{f_{td}(x); where\ x \in e3Dset|0 < d \leq i\})$ (eq. 4)
$M_D(ftriad, i) = Card(\{f_{td}(x)|where\ x \in f3Dset\ and\ (i-1) < d \leq (i)\})$ (eq. 5)
$M_c(ftriad, i) = Card(\{f_{td}(x)|where\ x \in f3Dset\ and\ 0 < d \leq i\})$ (eq. 6)

Then, we define the frequencies (F for triads, G for extended-triads and H for feature-triads) as:

$F(triad, i) = N(triad, i)/\sum_{j=1}^{30} N(triad, j)$ (eq. 7)
$G(etriad, i) = L(etriad, i)/\sum_{j=1}^{30} L(etriad, j)$ (eq. 8)
$H(ftriad, i) = M(ftriad, i)/\sum_{j=1}^{30} M(ftriad, j)$ (eq. 9)

Where N can be $N_D$ or $N_c$, L can be $L_D$ or $L_c$ , and M can be $M_D$ or $M_c$, depending on the approach to group the triads. This definition forces us to consider independent the groups obtained by cut-offs, instead of using the ratios with respect to the limit at 30 Å. We tested in artificial data that this approach preserves the curve of the statistical potential similar to the classical definition by bins of distances, but it's less affected by the scarcity of data.

To define a reference-state for the statistical potential, we require two more frequencies, one for $N_D$ and another for $N_c$ , using the **total number of triads** in the database (***triads***):

$O(i) = \sum_{triads} N(triad, i) \,/\, \sum_{j=1}^{30} \sum_{triads} N(triad, j)$ (eq. 10)

Where $triad \in triads$, and it's easy to proof that:

$$\sum_{j=1}^{30} \sum_{triads} N(triad, j) = \sum_{j=1}^{30} \sum_{ftriads} M(ftriad, j) = \sum_{j=1}^{30} \sum_{etriads} L(etriad, j)$$
(eq.11)

Where ***ftriads*** is the set of feature-triads and ***etriads*** the set of extended-triads, with $ftriad \in ftriads$ and $etriad \in etriads$. Using these definitions and following previous works [13], we define the potentials E3DC, ES3DC and PAIR per *triad* and distance *d*, using the round value of d (i.e. k), as follows:

$k = 1 + int(d-1)$
$PAIR(triad, d) = K_B T log_e(O(k)) - K_B T log_e(F(triad, k))$ (eq.12)
$ES3DC(etriad, d) = K_B T log_e\big(H(ftriad, k)\big) - K_B T log_e(G(etriad, k))$ (eq. 13)
$E3DC(ftriad, d) = -K_B T log_e\big(O(k)\big) + K_B T log_e(H(ftriad, k))$ (eq. 14)

Where, F, G, H and O are frequencies calculated by bins or using cut-offs.

The total potential of an interaction is calculated as the sum of the corresponding potential of all triads, feature-triads and extended-triads at distances shorter than 30Å.

We use each potential to **score the quality** (or potentiality) of the interaction. Let be I, E and D the sets defined respectively as the set of all triads, extended-triads and feature-triads with their associated distances (d), amino-acid residue number (p) and dinucleotide position (q) in the binary interaction of a TF-DNA structure. Therefore, we define the **energy-based** ***scores*** (as they are based on total potentials) as:

$$PAIR = \sum_{(triad,d,p,q)\in I} PAIR(triad,d) \quad \text{(eq. 15)}$$
$$ES3DC = \sum_{(etriad,d,p,q)\in E} ES3DC(etriad,d) \quad \text{(eq. 16)}$$
$$E3DC = \sum_{(ftriad,d,p,q)\in D} E3DC(ftriad,d) \quad \text{(eq. 17)}$$

Similarly, we also use the potentials defined for *triads, ftriads* and *etriads* as *scores* of the quality of a single interaction between one dinucleotide and one amino-acid residue. Then, the potential ES3DC of an extended-triad associated with distance d can be rewritten as:

$$ES3DC(etriad,d) = E3DC(ftriad,d) + K_B T log_e\big(O(k)\big) - K_B T log_e(G(etriad,k)) \quad \text{(eq. 18)}$$

Where $k = 1 + int(d-1)$. We then define two scoring terms, one distance independent ($ES3DC_{di}$) and another distance dependent ($ES3DC_{dd}$), as follows:

$$ES3DC_{di}(etriad) = K_B T log_e\big(\sum_{j=1}^{30} L(etriad,j)/\sum_{j=1}^{30}\sum_{triads} N(triad,j)\big) \quad \text{(eq.19)}$$
$$ES3DC_{dd}(etriad,d) = -K_B T log_e(L\,(etriad,k)/\sum_{triads} N(triad,k)) \quad \text{(eq. 20)}$$

Using eq. 8, eq. 19 and eq. 20, we rewrite $ES3DC(etriad,d)$ as:

$$ES3DC(etriad,d) = E3DC(ftriad,d) + ES3DC_{di}(etriad) + ES3DC_{dd}(etriad,d) \quad \text{(eq. 21)}$$

And the global energy-based *scores:*

$$ES3DC_{dd} = \sum_{(etriad,d,p,q)\in E} ES3DC_{dd}(etriad,d)$$
$$ES3DC_{di} = \sum_{(etriad,d,p,q)\in E} ES3DC_{di}(etriad) \quad \text{(eq.22)}$$

We consider $ES3DC_{dd}$ to evaluate the quality of an interaction because it is more specific than PAIR, as it uses extended-triads and distances, and it is also highly sensible, because it is normalized over the total of triads at a given distance with independence of the features. However, we have to note that this can only be considered as a scoring of quality, because the complete statistical potential requires the other terms E3DC and $ES3DC_{di}$ to complete ES3DC in equation 21.

Finally, due to the scarcity of data the curves of statistical potentials may be jagged. Therefore, we use a sliding window of approximately W samples defined by distances (by bins or cut-off) to **smooth the potential curves**. Let be $Scr(d)$ a distance-dependent score, as defined previously, then we define the smoothed score, $Scr_{smooth}$, as:

$$Scr_{smooth}(d) = \frac{1}{size_{samples}} \sum_{k=k_{min}}^{k_{max}} Scr(k)$$

(eq.23)

Where $k_{max}$ and $k_{min}$ are defined as:

$$k_{max} = \min(30;\ 1 + int(d-1) + int(W/2))$$
$$k_{min} = \max(0;\ 1 + int(d-1) - int(W/2))$$

Where, $size_{samples}$ is the total of bins between $k_{min}$ and $k_{max}$ with defined score ($Scr$), which is around W (i.e. $size_{samples} \approx W$)

### 5. Z-scores

We have to note that frequencies are always smaller than 1, consequently their logarithm is smaller than 0. However, statistical potentials are not necessarily negative, because by definition the use of a reference state is required, which is obtained by the sum of all triads at a given distance (also *ftriad* or *etriad*, depending on the potential of interest). Consequently, the comparison with a reference state can change the sign of the final sum of terms (for example, PAIR and ES3DC can be negative while E3DC may be positive). The variability of signs of the potentials affects the criterion of quality of the scores and they may become unclear. Nevertheless, indistinctly of the sign, the best interaction between an amino-acid and a dinucleotide is produced at the distance where PAIR and ES3DC are minimum, because it implies the highest frequency of a *triad* (or *etriad*) with respect to all triads (or feature-triads). For example, $ES3DC_{dd}$ has a minimum for the highest frequency of an extended-triad with respect to all triads. We then define *z-scores* in order to follow a criterion that incorporates the sign to score the quality of the interaction between an amino-acid and a dinucleotide as a function of the distance. We wish that the z-score identifies simultaneously the best distance associated with a triad and the best pair formed by one amino-acid and one dinucleotide. Consequently, we construct a ***zscore* function** with any type of *score*, applying without loss of generality on an extended-triad (*etriad*) and an associated distance d, as:

$$zscore(score) = \frac{score(etriad,d) - \mu_A\big(score(etriad,d)\big)}{\sigma_A\big(score(etriad,d)\big)}$$

(eq. 24)

Where: A is the set of all amino-acid types (i.e. $Car\{A\} = 20$), and we use the classical functions of **average** ($\mu$) and **standard deviation** ($\sigma$), defined as:

$$\mu_A\big(score(etriad,d)\big) = \frac{1}{Car\{A\}} \sum_{r \in A} score\big(\epsilon_a(etriad,r)\big)$$

$$\sigma_A\big(score(etriad,d)\big) = \frac{1}{Car\{A\}} \sqrt{\sum_{r \in A} \Big(score(etriad,d) - \mu_A\big(score(etriad,d)\big)\Big)^2}$$

Notice that we use the function $\varepsilon_a(etriad, r)$ as above and that we use the term *score* instead of potential (P in previous section) to emphasize that this is used only as a criterion of the quality of the interaction.

It's easy to proof that this definition satisfies our requirements for the new function *zscore*: 1) the minimum value of *score* produces a negative value of *zscore* (and *vice-versa*, the maximum yields positive); 2) as the *zscore* compares the *score* of a particular amino-acid with all other, the *etriad* becomes ranked from minimum to maximum allowing us to select the best residue among amino-acids in the same position, depending on the quality criterion of the score. Finally, if we choose to smooth the curves dependent on distance, the *zscore* has to be calculated first with the unsmoothed scores and subsequently smoothed to avoid introducing biases.

### 6. Structural modeling of C2H2-ZF complexes

Given the sequence of a TF (named protein target) and a DNA binding site fragment (named DNA target), we obtain the structure of the complex by means of homology modelling using the program MODELLER [7]. First, we search potential templates of the protein target among the set of sequences with known structure in $PDB_{DNA}$ using BLAST [15]. We use the alignment obtained with *matcher* and the sequence and structure of each template to construct the models with MODELLER.

The DNA binding sequence of each testing 3-domain C2H2-ZF protein is formed by 9bp nucleotides, a set of 3bp bound by each individual domain. For the selection of the binding sequences associated with each finger domain we use the same sequences as in the B1H experiment [11]: 1) for the selection of F2, the finger sequences in F1 (N-tail domain) and F3 (C-tail domain) involved in the interface are RSDNLRA(F1) and RSANLVR (F3), respectively binding AAG and GAG; and 2) for the selection of F3, the finger sequences in F1 (N-tail domain) and F2 (inner domain) involved in the interface are RSDELTR (F1) and RSDNLRA (F2), respectively binding GCG and AAG.

The structure of Zif268 binding DNA is modelled with 23 different template structures to introduce structural variability, retrieved by codes: 1p47 (chain A), 1zaa (chain C), 1g2d(chain C), 1a1h(chain A), 1a1g(chain A), 1a1i(chain A), 1a1j(chain A), 1a1k(chain A), 1a1l(chain A), 1aay(chain A), 1jk1(chain A), 1jk2(chain A), 2kmk(chain A), 2wbu(chain A), 4r2a(chain A), 4r2c(chain A), 4r2d(chain A), 5ke6(chain A), 5ke7(chain A), 5ke8(chain A), 5ke9(chain A), 5kea(chain A) and 5keb(chain A) . We compare the sequence of Zif268 used in the experiment with the templates by a sequence alignment with CLUSTALW [16]. We identify as WT the original sequence, and as F2 and F3 the sequences used for the selection of F2 and F3 binding sequences, labeling by "X" the amino-acids that are modified. This alignment is shown here for 1p47 used as template, highlighting in bold and red the attention of the specific binding sequence of each finger (blue box for F1, yellow box for F2 and green box for F3):

```
F2          GTERPYACPVESCDRRFSRSDNLRAHIRIHTGQKPFQCRICMRNFSXXXXLXXHIRTHTG
F3          GTERPYACPVESCDRRFSRSDELTRHIRIHTGQKPFQCRICMRNFSRSDNLRAHIRTHTG
WT          GTERPYACPVESCDRRFSRSDELTRHIRIHTGQKPFQCRICMRNFSRSDNLRAHIRTHTG
1p47_A      --ERPYACPVESCDRRFSRSDELTRHIRIHTGQKPFQCRICMRNFSRSDHLTTHIRTHTG
              ****************    *  *********************    *  *******

F2          EKPFACDICGRKFARSANLVRHTKIHLRGS
F3          EKPFACDICGRKFAXXXXLXXHTKIHLRGS
WT          EKPFACDICGRKFARSANLVRHTKIHLRGS
1p47_A      EKPFACDICGRKFARSDERKRHTKIHLRQ-
            **************       ******
```

After modeling the structure of Zif268, we complete the complex by modeling the structure of the DNA binding sequence. However, each template has DNA sequences of different length that do not correspond with the DNA used in the experiment. The full DNA sequence of the experiment is longer (29bp) than the binding (9bp), which is shown next, embedded in positions 11 to 19 (nucleotides highlighted), labelling by "N" those under test. Two DNA sequences are considered depending on the experiment, one for the selection of F2 and another for F3:

```
F2:    5'- GCGGCCGCAAGAGNNNAAGTAACGAATTC - 3'
F3:    5'- GCGGCCGCAANNNAAGGCGTAACGAATTC - 3'
```

The structure of the full DNA sequence bound by Zif268 is obtained with the program X3DNA[4] by modifying the DNA structure in the complex. First, we locate the 9bp binding region in the structure of the template and identify the positions at 5' and 3' (*first* and *last*). Next, we construct two frames of B-DNA structure with the lengths required to extend the template at 5' and 3' up to 29 nucleotides. The lengths in both sides depend on the location of the 9bp binding sequence and the DNA sequence of the experiment. Then, we use 3DNA/DSSR to perform a least-squares fitting that locates each base reference frame in the *first* and *last* positions. Finally, the structure of DNA is completed with the right sequence. We model the DNA structure by substitution of the nucleotides in each model with the corresponding nucleotides of the target. We use the program X3DNA to substitute the nucleotides of one strand and automatically model its corresponding pair.

We also model several structures with the complex of Zif268 binding a non-specific DNA region using the same approach. These structures are used as non-binding examples. The non-binding sequence is taken randomly by selecting a region of the sequence of the weak promoter GAL1, constructing 10 DNA fragments of 29bp for each known binding. A potential binding test in this region, which is part of the B1H experiment, may well represent the background. The forward weak promoter sequence of GAL1 is formed by 118bp that are shown here:

```
5'- GAGATTAAGGAGCAGAAGGGGTGACAGCCCTCCGAAGGAAGA
    GAGATTAAGCTCTCCTCCGTGCGTCCTCGTCTTCACCGGTCG
    CGTTCCTGAAACGCAGATGTGCCTCGCGCCGCACTGCTCCG - 3'
```

### 7. Use of experimental TF-DNA binding to calculate statistical potentials

One of the main problems to obtain statistical potentials for all families and folds of TFs is the scarcity of known interactions. We need to enlarge the number of interacting triads. Therefore, here we propose to use the experimental knowledge of TF-DNA interactions to derive interacting triads without requiring the complete knowledge of TF-DNA complex structures. We then use the sets of derived triads associated with distances to calculate the statistical potentials.

Let be a TF with experimentally known interactions with several DNA sequences. We define a mapping function (*map*$_D$) between the fragment of the DNA sequence from the structure ($S_{DNA}$ ), and each positive binding sequence ( $S_{exp}$), as follows:

$$n_m, q_n = map_D(v_m, q_v)$$
(eq.25)

Where $n_m$ is a dinucleotide, with $n_m \in S_{exp}, v_m$ is also dinucleotide, with $v_m \in S_{DNA}$, $m$ is the position of the dinucleotide in the alignment between $S_{exp}$ and $S_{DNA}$, $q_v$ is the position of the dinucleotide $v_m$ in $S_{DNA}$, $q_n$ is the position of the dinucleotide $n_m$ in $S_{exp}$, and the position of each dinucleotide is defined (i.e. equal to) the position of the first nucleotide in the DNA sequence. For the sake of simplicity, when the length of $S_{DNA}$ is the same as $S_{exp}$ and the position in the alignment, $m$, coincides with $q_v$ and $q_n$ (i.e. $m = q_n = q_v$ ), we write:

$$n_m = \widehat{map}_D(v_m)$$
(eq.26)

We can define a set of mapping functions with the alignments of all the DNA sequences extracted from the structures of TF-DNA complexes in $PDB_{DNA}$ that are aligned with positive binding sequences. Then, we use the dinucleotide substitution function (as defined in section 3), $\eta_v(etriad, n)$, of an extended-triad, *etriad*, containing a nucleotide $v$ which is substituted by *n* in the dinucleotide, to generate more extended-triads.

There are other TFs for which the structure is not known but can be modelled. This implies that the sequence of the TF can be aligned with sufficient percentage of identical residues to ensure its modeling. Then, we define another mapping (*map*$_P$) between the protein sequence of the TF with known experimental data on DNA binding and the TF sequence of a known structure (template), with:

$$r_m, p_r = map_P(a_m, p_a)$$
(eq.27)

Where, $a_m$ is an amino-acid residue in the sequence of a template and $r_m$ is the amino-acid in the sequence of a TF with experimental data, in position $m$ of the alignment of both TF sequences that correspond with positions $p_r$ and $p_a$ for $r_m$ and $a_m$, respectively. Also, for the sake of simplicity, if the position in the alignment, $m$, coincides with $p_a$ and $p_r$ (i.e. $m = p_r = p_a$ ) we write:

$$r_m = \widehat{map}_P(a_m)$$
(eq.28)

We define the set of all mapping functions with all the alignments between the sequences of TFs with experimental annotation and the TFs with known structure (complexed with DNA). Also, we define $etriads(a, v)$ as the set of extended-triads, extracted from the 3Dset, containing amino-acid residue $a \in A$ (the set of 20 amino-acids) and dinucleotide $v \in \Lambda = \{A, C, G, T\} \times \{A, C, G, T\}$.

We remind the definition of $\varepsilon_a(etriad, r)$ as the function that substitutes amino-acid residue "a" of an extended-triad, *etriad*, by the amino-acid residue "r". With all these definitions, we increase the set of extended-triads of the e*3Dset* to *e3Dset'*, using all the mapping functions *map<sub>D</sub>* (simplified as $\widehat{map}_D$) and *map<sub>P</sub>* (simplified as $\widehat{map}_P$). We use the simplified maps without loss of generality, as all sequences and alignments can be renumbered. Both mappings, $\widehat{map}_D$ and $\widehat{map}_P$, are respectively defined with: 1) the alignments of the DNA sequences extracted from the structures in the PDB aligned with positive binding sequences; and 2) the TFs that can be modelled using the alignment with their structural templates (the set is defined as $PBM_{3Dset}$). The new set of extended-triads, *e3Dset'*, is defined as:

$$e3Dset' = \left\{ (etriad, d, p, q) \middle| \begin{array}{c} with\ etriad = \eta_v\left(\varepsilon_a\left(x, \widehat{map}_P(a)\right), \widehat{map}_D(v)\right);\ x \in etriads(a, v); \\ x\ associated\ with\ d, p\ and\ q;\ \forall a \in A;\ \forall v \in \Lambda;\ \forall\ \widehat{map}_D, \widehat{map}_p \in PBM_{3Dset} \end{array} \right\}$$

(eq. 29)

And we recalculate $L_D$ and $L_C$ in equations eq.3 and eq.4 as:

$L_D(etriad, i) = Card(\{f_{td}(x) | where\ x \in e3Dset'and\ (i-1) < d \leq (i)\})$ (eq. 30)
$L_c(etriad, i) = Card(\{f_{td}(x) | where\ x \in e3Dset'and\ 0 < d \leq i\})$ (eq. 31)

Similar approach is also taken for the sets of *triads* and *ftriads*, modifying accordingly the corresponding functions to substitute amino-acid and dinucleotide residues of a triad or a featured-triad.

We proceed similarly with B1H experiments on C2H2-ZF family. For each finger (F2 and F3) and combination of 3bp nucleotides, we collect all protein sequences producing significant binding signal in the B1H experiment. We use the modelled structures of the DNA testing sequence of 29bp with different templates and introduce the mappings for the DNA sequence and the modified residues in F2 or F3 from the multiple sequence alignment. For the DNA sequence the mapping is on the 3bp modified nucleotides, affecting 4 dinucleotides, while for the protein sequence the mapping affects 6 amino-acids, both mappings being different for F2 and F3 selections. This is, for the DNA sequence the mapping is $n_m = \widehat{map}_D(v_m)$, where $n_m$ is a dinucleotide of the 3bp under test, $v_m$ is a dinucleotide of the 29bp of the modelled template, and the position, $m$, is between 11 and 19 (11-13 for F3 and 14-16 for F2). For the substitution of residues of Zif268 in the interface we require a mapping of the native sequences RSANLVR (F3) and RSDNLRA (F2), for all selections of the binding finger. This mapping is $(r_m, p_r) = map_P(a_m, p_a)$ where $m$ is the position in the alignment (residues 47-53 for F2 and 75-81 for F3), $p_r = m$, the position in the template, $p_a$, depends on the template used to model Zif268 and $a_m$ is the corresponding residue in the template in position $m$ of the alignment. From the template structure we extract the contacts (*triads, etriads* and *ftriads*) between amino-acids and dinucleotides and generate the statistical potentials. However, this introduces a bias by overestimating the constant amino-acids and nucleotides that have not been modified in the experiment. Therefore, only the triads affecting the amino-acids and nucleotides under test are considered to generate the potentials. This is, we only consider the contacts between the 6 amino-acids labelled by "X" and dinucleotides containing one of the 3bp labelled by "N" to generate two potentials, one for F2 and another for F3 finger positions. We restrict each set of Zif268 sequences to those with highest signal of the B1H binding experiment in order to obtain potentials more specific or associated with the strongest binding. We define three thresholds based on the affinity percentile of a sequence: 1) higher than 90%; 2) higher than 75%; and 3) higher than 50%. To calculate the affinity percentile of a sequence we follow the same definition as the authors[11]. Each sequence, $seq$ in its corresponding domain, has a logged and normalized frequency of its observation ($p_{seq}$). Hence, the affinity percentile is defined as the sum of all other frequencies lower or equal to the frequency of the sequence (e.g. an affinity percentile of 90% implies that the sequence is on the tail with highest number of observations, in the top 10%). However, the number of selected sequences may be too different between experiments (i.e. the 3 nucleotides of the binding "TGA" may have 30 sequences with affinity percentile higher than 90, while AAA has only 2), which produces the opposite bias on the expected potential. To avoid a bias on the number of sequences selected, we force to have around 500 sequences for all trinucleotide-binding experiments, by repeating as many times as we need each sequence (e.g. if only 2 sequences are selected for AAA and they are equally representative, we should repeat 250 times each). Each sequence is repeated in proportion to the number of observations. This is, for a sequence with observation $p_{seq}$, it is repeated $500 \times \frac{p_{seq}}{\sum_{k \in A_{90}} p_k}$, where $A_{90}$ is the set of sequences with affinity percentile higher than 90 (or we use $A_{50}$ for affinity percentile higher than 50, etcetera). As a consequence of the approach, the contacts derived from the B1H experiment are limited to relatively short distances (the largest contacts are around 15-20Å). However, we note that we also use contacts extracted from other structures of the C2H2-ZF family in the PDB and from the use of PBM experiments, covering larger distances up to 30 Å.

### 8. Scoring TF-DNA binding with structure

#### a. Scores of single domain structures

Given the structure of a protein-DNA complex, either experimentally obtained (i.e. from crystallography and identified by a PDB code) or modelled, we define several scores of the interaction based on statistical potentials. First, we calculate the interface of the interaction and extract all triads, extended-triads and feature-triads associated with distances shorter than 30Å. Then, the score of the interaction is defined as the sum of the scores (i.e. potential) of all triads with their associated distances (or extended-triads or feature-triads, depending on the type of score). The same approach is applied for z-scores. Let $Scr$ be a potential or a z-score and let C be the set of triads (extended-triads or feature-triads, depending on the definition of $Scr$) and their associated distances (d), amino-acid residue number (p) and dinucleotide position (q). The score of the interaction is defined as:

$$score_{Scr} = \sum_{(triad,d,p,q)\in C} Scr(f_{td}(triad, d, p, q))$$

(eq.32)

We can obtain the score of a TF without knowing the structure of the TF-DNA binary complex if it can be modelled. We use the structure of a template to generate the set of triads and the mapping of amino-acids derived from a sequence alignment, $map_P$ as defined in section 7 (eq.28), between the TF sequence and the sequence of the template. We also need the mapping of dinucleotides between the DNA sequence we wish to model and the DNA sequence in the template interface. Instead of modeling the structure of the TF-DNA complex, we modify the scores by applying the substitution of the corresponding amino-acids, using the functions defined as in section 3, $\varepsilon_a(triad, r)$ and $\eta_v(triad, n)$, and the mappings $map_P$ (in eq. 28) and $map_D$ (in eq. 27) between the templates and the sequences of TF and DNA, respectively. Here, instead of using simplified mappings ($\widehat{map}_D$ and $\widehat{map}_P$), we generalize the formula by defining special functions, $f_1$ and $f_2$, to extract the dinucleotide or amino-acid positions in the DNA or protein sequences:

$$f_1(r, q) = r$$
$$f_2(r, q) = q$$

(eq. 33)

Where $r$ is either a dinucleotide or an amino-acid residue, and $q$ is a position of a dinucleotide or an amino-acid, respectively for DNA or protein sequences. Then we calculate the score of the interaction as:

$$score_{Scr} = \sum_{(triad,d,p,q)\in C} Scr(\eta_v(\varepsilon_a(triad, f_1(map_P(a, p))), f_1(map_D(v, q))), d)$$

(eq.34)

#### b. Multiple domain TFs

There are TF structures with more than one domain, such as the particular case of the C2H2-ZF family, where the TF has several domains like F2 (internal) and two more domains, one at the N-tail (F1) and another C-tail (F3). However, we apply two potentials for all domains, one obtained with B1H data on finger domains in F2 and another for F3. For example, we use the statistical potential ZES3DC calculated with variant sequences in F2 domain (**ZES3DCF2**) or in F3 (**ZES3DCF3**).

### 9. Construction of PWMs using Zif268 structure models.

Given the modelled structure of Zif268-DNA complex, we obtain the PWM by means of statistical potentials using scores or zscores (we use the zscore of $ES3DC_{dd}$ by default as example). We focus on the specific nucleotides for three continuous fingers (9 bases) covering the whole binding site in sliding windows. We collect the set of triads, extended-triads and feature-triads with their associated distances between protein and DNA and the associated amino-acid and dinucleotide positions (i.e. $C_k$, $eC_k, fC_k$, respectively), where the dinucleotide in the triad belongs in 9 nucleotide overlapping fragments (we name them $DNA_{F_1}$ , $DNA_{F_2}$ etc.). We remind the substitution function $\eta_v(etriad, n)$, to substitute the dinucleotide $v$ of an $etriad$ by the dinucleotide $n$. Similar functions are defined to substitute the dinucleotide in triads and feature-triads (these are only affected in the change of the nitrogenous bases).

Then, for each fragment $DNA_{F_k}$ , with k=1,N-8 and the binding site of length N, we obtain a test set with all possible DNA sequences (i.e. a total of $4^9$). We define two mappings, $map_{DNA_{F_k}}$ and $map_{seq}$ , respectively for the native fragment sequence of the model and any sequence $seq$ in the test set, both between sequence position and dinucleotides (i.e. $map_{seq}(j) = \omega_j\omega_{j+1}$ with $\omega_j$ nucleotide in position $j$ of $seq$). We calculate the score of any sequence of the test set with the triads, extended-triads and feature-triads, using the associated distances, residue number and dinucleotide positions from the complex structure (i.e. triads, extended-triads and feature-triads as $C_k$, $eC_k, fC_k$, respectively) and using the corresponding substitution function (e.g. $\eta_v(etriad, n)$). Let $ZES3DC_{dd}$ be the z-score of $ES3DC_{dd}$ and assume we apply it on extended-triads without loss of generality, then the score of a sequence $seq$ is:

$$score_{seq} = \sum_{(etriad,d,p,q)\in eC_k} ZES3DC_{dd}(\eta_v(etriad, n), d, p, q) \qquad \text{(eq. 35)}$$

Where, for each $(etriad, d, p, q) \in eC_k$, $etriad, d, v$ and $n$ are calculated using the functions as defined in section 3 and the mappings defined above:

$$\begin{aligned} etriad &= f_t(etriad, d, p, q) \\ d &= f_d(etriad, d, p, q) \\ q &= f_n(etriad, d, p, q) \\ v &= map_{DNA_k}(q) \\ n &= map_{seq}(q) \end{aligned}$$

We normalize the scores between 0 and 1, using the total set of scores obtained with all generated sequences (i.e. set $\{score_{seq}\}$ ) and modifying the sign accordingly:

$$normal(score_{seq}) = \frac{score_{seq} - \min(\{score_{seq}\})}{\max(\{score_{seq}\}) - \min(\{score_{seq}\})} \qquad \text{(eq.36)}$$

Where we have assumed that the best score is already the maximum. For example, for $ZES3DC_{dd}$ we need to multiply the score by -1. Then, we rank the normalized scores and select only the DNA sequences producing the top scores over a **cut-off threshold** (often 0.9). This produces an alignment from which we calculate the PWM (i.e. frequencies of nucleotides in each position)

#### 10. Score per Nucleotide: profiles of a DNA binding site.

We define a **nucleotide profile** as a function on $\mathbb{R}$, $f: \mathbb{N} \longrightarrow \mathbb{R}$, of the nucleotide position in a DNA sequence. Here, we define **score-nucleotide profiles** when the function is obtained with the scores and z-scores. Given a TF-DNA complex structure, **3D-TF**, and a distance dependent score, $score$, obtained with statistical potentials (e.g. the smoothed z-score of $ES3DC_{dd}$). Let be $j$ a nucleotide position of the DNA sequence in the complex, we define a new score, $Scr_{score}$ in $j$, as:

$$Scr_{score}(j) = \sum_{(etriad,d,p,q) \in eC_j} score(etriad, d)$$

(eq.37)

Here we use extended-triads without loss of generality, although depending on the statistical potential we can use triads or feature triads instead. The set $eC_j$ is the set of *etriads* with all associated distances (i.e. any $d$), and amino-acid residue numbers (i.e. any $p$), where dinucleotide in position $q$ implies that $j$ is the position of one of the nucleotides in the dinucleotide at $q$ (i.e. $q = j \; or \; q = (j-1)$). This new score can be normalized by considering the contribution of the nucleotide to the total score or as the percentage of the contribution of all nucleotides of the DNA sequence. Assuming that the score is the smoothed z-score of $ES3DC_{dd}$ and without loss of generality, the normalized nucleotide profile is:

$$Normal(j) = \frac{\sum_{(etriad,d,p,q) \in eC_j} \big(zscore(ES3DC_{dd}(etriad,d))\big)_{smooth}}{\sum_{j=1}^{length} \sum_{(etriad,d,p,q) \in eC_j} \big(zscore(ES3DC_{dd}(etriad,d))\big)_{smooth}}$$

(eq. 38)

or

$$Normal(j) = \frac{\sum_{(etriad,d,p,q) \in eC_j} \big(zscore(ES3DC_{dd}(etriad,d))\big)_{smooth}}{factor_j \times ZES3DC_{dd}}$$

(eq. 39)

Where $factor_j = 2$ for all $j$ but the extremes at 5' and 3', in which is 1, and from equation 37 we write $ZES3DC_{dd}$ as:

$$ZES3DC_{dd} = \sum_{(etriad,d,p,q)\in E} \Big(zscore\big(ES3DC_{dd}(etriad,d)\big)\Big)_{smooth}$$

(eq. 40)

$E$ is the set of extended-triads (with their associated distances, amino-acid numbers and dinucleotide positions) and $length$ is the length of the DNA sequence. The factor of 2 in equation 39 is produced by the fact that each nucleotide is counted twice in $eC_j$, with the exception of the extreme positions in 5' and 3' where the nucleotides are only reckoned one time.

The curve of $Scr_{score}(j)$ (raw or normalized) along the positions in the DNA sequence is defined as the **nucleotide profile based on 3D-TF** for the (raw or normalized) potential defined in $score$ (e.g. the smoothed z-score of *ES3DC<sub>dd</sub>*). Hence, profiles defined upon scores derived from statistical potentials are dependent on the structure of the TF-DNA interaction complex. We have to note that, if the structure of this complex has been modelled, several models may be considered (i.e. we define the set of models of TF-DNA as $MDL_{TF}$). Besides, some models may be obtained using different templates, implying that these models introduce a relevant variability on the conformational space of the TF-DNA interaction. Consequently, several nucleotide profiles of scores are accumulated for the same DNA sequence. We then calculate the average and standard deviation of $Scr_{score}(j)$ for all positions $j$ along the DNA sequence with the nucleotide profiles of score based on each model structure $m$ of the $MDL_{TF}$ set (i.e. described as $\big(Scr_{score}(j)\big)_m$ ), using following equations:

$$\langle Scr_{score}(j)\rangle = \frac{1}{Card(MDL_{TF})} \sum_{m\in MDL_{TF}} \big(Scr_{score}(j)\big)_m$$

$$RMSD\big(Scr_{score}(j)\big) = \sqrt{\frac{\sum_{m\in MDL_{TF}} \Big(\big(Scr_{score}(j)\big)_m - \langle Scr_{score}(j)\rangle\Big)^2}{Card(MDL_{TF})}}$$

(eq. 41)

Equations in 41 describe two new nucleotide profile functions. The average function is defined as the **nucleotide profile of $score$** (e.g. the smoothed z-score of *ES3DC<sub>dd</sub>*) and the RMSD defines its margins of error or variability.

The nucleotide profile can also be calculated with other "scores" different than those derived by statistical potentials, for example by the enrichment or the number of times that nucleotide *j* is selected by several PWMs assigned to a TF. This extends the definition of score nucleotide profiles to other scores different than those obtained with statistical potentials.

### 11. Construction of the experimental PWM

The experimental PWM of a Zif268 sequence obtained from B1H, with a specific hexamer fragment on F2 or F3, is calculated based on its affinities for different binding sites. The binding site is formed by thee nucleotides flanked by two fixed nucleotides (G and A for F2, and two A for F3). All binding sites targeted by a specific hexamer-fragment with affinity higher than a threshold are stored and gapless aligned to construct the PWM (e.g. the top 10% threshold uses all DNA-bound sequences with affinity percentile higher than 90%, while for a threshold of 100% we use all detected sites with any not null affinity percentile). We construct experimental PWMs for top 10%, top 25%, top 50% and for all targeted sites. These experimental PWMs are also named hexamer-specific PWMs, to distinguish from PWMs obtained with other experiments or with a different approach.

**Figure S1**

**Figure S1**

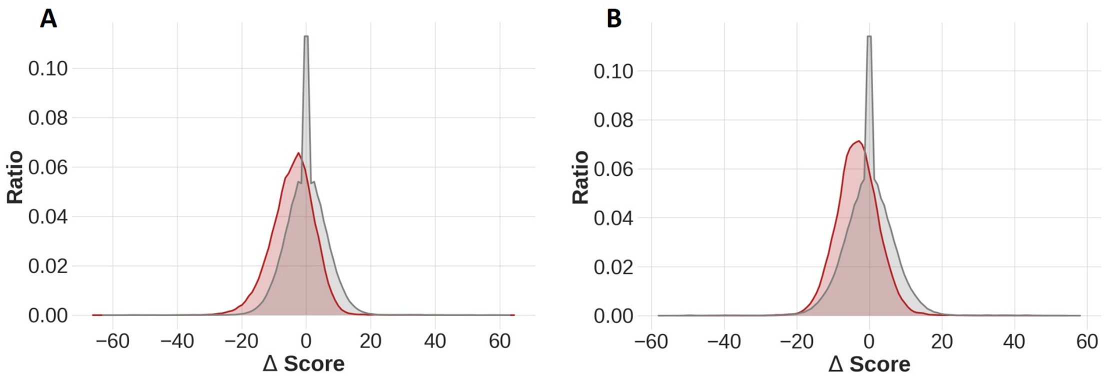

**Figure S2**

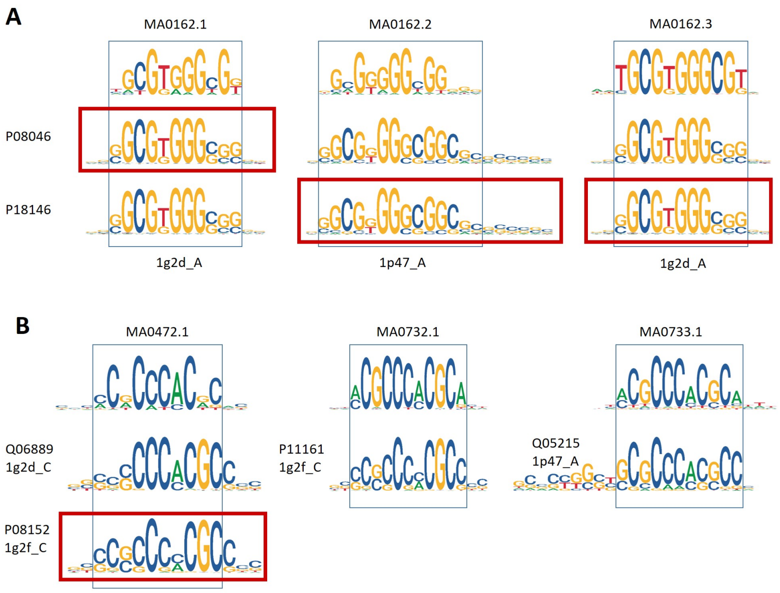
A
MA0162.1
MA0162.2
MA0162.3
P08046
P18146
1g2d_A
1p47_A
1g2d_A
B
MA0472.1
MA0732.1
MA0733.1
Q06889
1g2d_C
P11161
1g2f_C
Q05215
1p47_A
P08152
1g2f_C

**Figure S3**

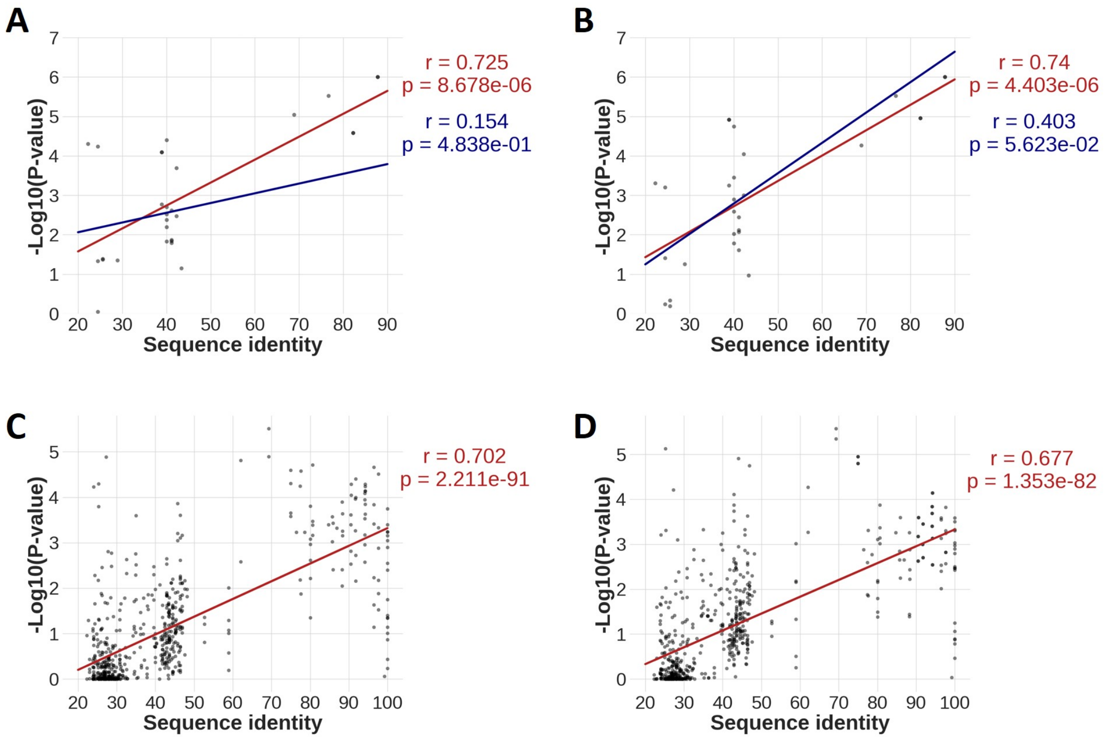
A
r = 0.725
p = 8.678e-06
r = 0.154
p = 4.838e-01
-Log10(P-value)
Sequence identity
B
r = 0.74
p = 4.403e-06
r = 0.403
p = 5.623e-02
-Log10(P-value)
Sequence identity
C
r = 0.702
p = 2.211e-91
-Log10(P-value)
Sequence identity
D
r = 0.677
p = 1.353e-82
-Log10(P-value)
Sequence identity

### Figure S4

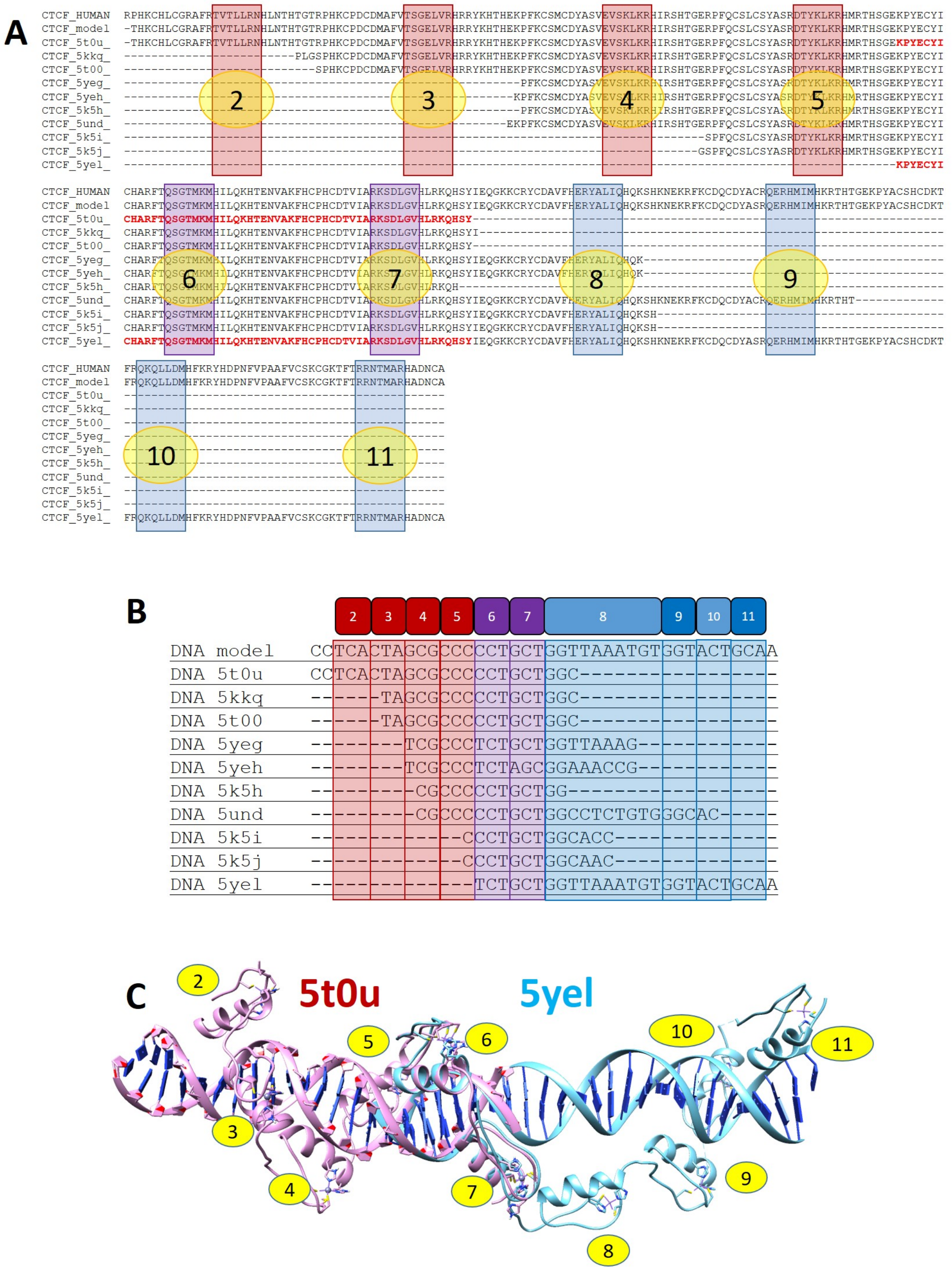
A
CTCF_HUMAN
CTCF_model
CTCF_5t0u_
CTCF_5kkq_
CTCF_5t00_
CTCF_5yeg_
CTCF_5yeh_
CTCF_5k5h_
CTCF_5und_
CTCF_5k5i_
CTCF_5k5j_
CTCF_5yel_
B
2 3 4 5 6 7 8 9 10 11
DNA model CCTCACTAGCGCCCCCTGCTGGTTAAATGTGGTACTGCAA
DNA 5t0u CCTCACTAGCGCCCCCTGCTGGC------------------
DNA 5kkq -----TAGCGCCCCCTGCTGGC------------------
DNA 5t00 -----TAGCGCCCCCTGCTGGC------------------
DNA 5yeg -------TCGCCCTCTGCTGGTTAAAG-------------
DNA 5yeh -------TCGCCCTCTAGCGGAAACCG-------------
DNA 5k5h --------CGCCCCCTGCTGG-------------------
DNA 5und --------CGCCCCCTGCTGGCCTCTGTGGGCAC------
DNA 5k5i ------------CCCTGCTGGCACC---------------
DNA 5k5j ------------CCCTGCTGGCAACC--------------
DNA 5yel --------------TCTGCTGGTTAAATGTGGTACTGCAA
C
5t0u
5yel

**Figure S5**

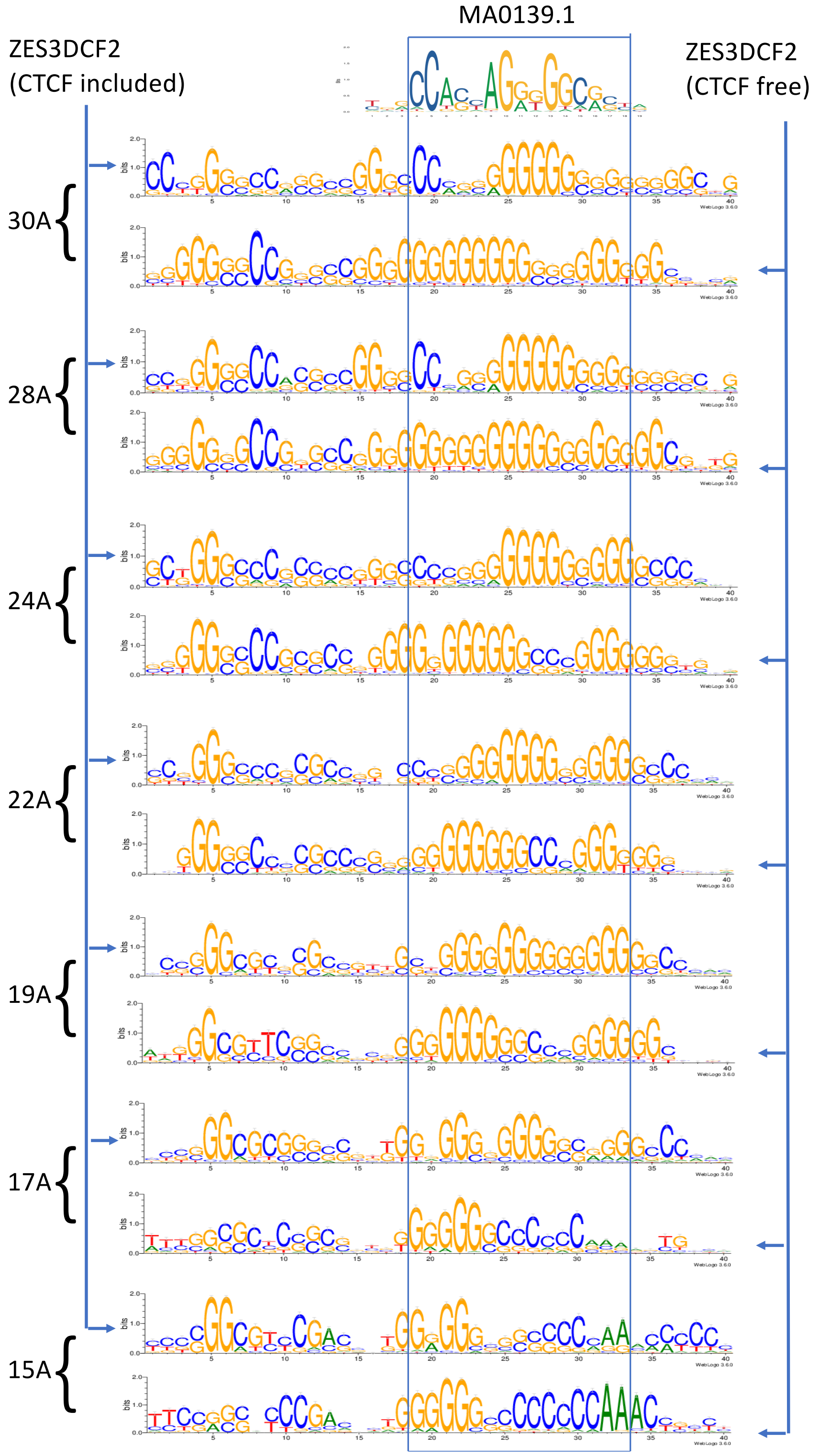
MA0139.1
ZES3DCF2
(CTCF included)
ZES3DCF2
(CTCF free)
30A
28A
24A
22A
19A
17A
15A
